# Evaluation of the Efficacy of Omega-3 Fatty Acids Formulations in the Lipopolysaccharide-Induced Acute Lung Inflammation Rat Model

**DOI:** 10.1101/2021.11.23.469790

**Authors:** Chandrashekhar Kocherlakota, Nagaraju Banda, Arjun Narala, Srinath Akula, Kumar S.D. Kothapalli, J.T. Brenna

## Abstract

Many current treatment options for lung inflammation and thrombosis come with unwanted side effects. The natural omega-3 fatty acids (O3FA) are generally anti-inflammatory and antithrombotic. The O3FA are always administered orally and occasionally by intravenous (IV) infusion. The main goal of this study is to determine if O3FA administered by inhalation of a nebulized formulation mitigates LPS-induced acute lung inflammation in male Wistar rats. Inflammation was triggered by intraperitoneal injection of LPS once a day for 14 days. One hour later, rats received nebulized treatments consisting of egg lecithin emulsified O3, budesonide and Montelukast, and blends of O3 and melatonin or Montelukast or Cannabidiol; O3 was in the form of free fatty acids for all groups except one group with ethyl esters. Lung histology and cytokines were determined in n=3 rats per group at day 8 and day 15. All groups had alveolar histiocytosis severity scores half or less than that of the disease control (Cd) treated with LPS and saline only inhalation. IL-6, TNF-α, TGF-β, and IL-10 were attenuated in all O3 groups. IL-1β was attenuated in most but not all O3 groups. O3 administered as ethyl ester was overall most effective in mitigating LPS effects. No evidence of lipid pneumonia or other chronic distress was observed. These preclinical data suggest that O3FA formulations should be further investigated as treatments in lung inflammation and thrombosis related lung disorders, including asthma, chronic obstructive pulmonary disease, lung cancer and acute respiratory distress like COVID-19.

---

Pulmonary disorders, both acute or chronic such as acute respiratory distress syndrome (ARDS) [1], SARS-CoV-2 infections (causative agent of COVID-19) [2], asthma [3, 4], cystic fibrosis (CF) [5], chronic obstructive pulmonary disease (COPD) [6], interstitial lung disease (ILD) [7] or other lung disorders [8, 9] are commonly accompanied by pulmonary inflammation. The global COVID-19 pandemic caused by the SARS-CoV-2 virus is an enigma in part because of its wide clinical spectrum ranging, from complete silence to mild to severe clinical conditions such as respiratory failure, sepsis, cytokine storm, thrombosis and multiorgan dysfunction syndromes (MODS) [10, 11]. Since the first COVID-19 cases emerged in Wuhan, China, about 222 countries and territories have been affected with over 250,895,264 positive diagnoses and 5,068,954 deaths worldwide as of November 8, 2021 [12]. Both inflammation and thrombosis are mediated by signaling molecules derived from highly unsaturated fatty acids (HUFA) or more precisely the relative mix of them present at any one time in cell membranes [11].

Omega-3 (ω3 or n-3) fatty acids (O3FA), especially HUFA docosahexaenoic acid (DHA, 22:6n-3), docosapentaenoic acid (DPA, 22:5n-3) and eicosapentaenoic acid (EPA, 20:5n-3), are natural healthy fats which inhibit formation of proinflammatory/prothrombotic lipid mediators eicosanoids and docosanoids, and independently serve as substrates for production of anti-inflammatory and inflammation-resolving resolvins and protectins [13, 14]. Omega-3 HUFA are found in sea food, marine oils (e.g. finfish, squid, and krill oils) and microalgae [15–17]. Omega-3 are ubiquitous in vertebrate tissue, acting as bioactive components of cell membrane phospholipids and anchoring proteins in cell membranes. Their modification by biosynthetic inhibition or receptor-mediated actions remains a prevailing strategy for developing valuable drug targets used over-the-counter and prescription drugs such as aspirin, non-steroidal anti-inflammatory drugs (ibuprofen and naproxen) and leukotriene receptor inhibitors (zafirlukast, montelukast, and zileuton) [11, 18–22]. Eicosanoids and docosanoids have wide-ranging functions in the body’s cardiovascular, pulmonary, neurological, immune, and endocrine systems [23–26].

Treatment options for lung inflammation include corticosteroid or glucocorticoid or leukotriene receptor antagonist or many other medications, such as budesonide, prednisone, methylprednisolone, hydrocortisone or montelukast. However, many of these medications come with unwanted side effects that add health risks, or cause physical discomfort [27–29]. Drug interactions between antimicrobial agents and corticosteroids leading to Cushing’s syndrome, adrenal suppression, weight gain, osteoporosis, and steroid accumulation have been reported [30, 31]. Similarly potential risk of adrenal insufficiency due to interaction of glucocorticoids with antiviral therapy are issues in SARS-CoV-2 patients [32]. O3FA are natural, dietary and also available as supplements. O3FA protect against several types of lung diseases such as COVID-19 [33], asthma [34], cystic fibrosis [35], ARDS [36], COPD [37], and non-small cell lung cancer (NSCLC) [38].

The O3FA are always administered systemically, primarily orally and less commonly by intravenous (IV) infusion. When administered orally they are provided primarily in four common forms: as ethyl esters (EE), as triacylglycerols (TAG), as phospholipids (PL) and as free fatty acids (FFA), also known as non-esterified fatty acids (NEFA) [39–41]. In foods, TAG and PL are the overwhelmingly predominant forms, with small amounts of FFA. Only small amounts of FA EE are present in humans, usually endogenously synthesized upon consumption of ethanol (alcohol) [42]. FA EE usually synthesized by industrial processes from the natural forms, primarily TAG, for further purification. When administered intravenously, omega-3 are primarily provided as TAG as an emulsion with smaller amounts in PL that may originate with emulsifying agents such as egg PL. FA EE may be administered IV as emulsions [43, 44].

In any of these forms, systemic administration results in rapid hydrolysis (“lipolysis”) of all forms, where the resulting liberated FFA enter normal biochemical pathways present to transport and distribute FA to the blood stream to perfuse all organs. Smaller amounts of O3FA are captured and re-esterified into the various lipid classes in cells. In the bloodstream, O3FA are rapidly taken up in an untargeted manner into all tissues thus distributing the oral or intravenous dose to all organs. Thus, only a fraction of any given dose will be incorporated into any particular tissue, such as the lung.

Bioactivity/efficacy of O3FA against pathologies depend on their concentrations in target tissue and more specifically target lipids of target tissue. For instance, efficacy against lung pathology depends on the specific concentration of O3FA in lung tissue and more specifically the concentration in PL present in cell membranes and possibly surfactant lipids. Because of the untargeted nature of systemic administration, any particular dose is less efficacious than an equivalent dose delivered directly to the target organ. More specifically, any particular dose will be less efficacious for treating lung pathology when administered systemically as one of the common forms compared to an equivalent dose administered directly to the lung.

We hypothesized that delivery of O3FA by inhalation would be efficacious against inflammatory sequelae induced experimentally by intraperitoneal injection of lipopolysaccharide (LPS) as a model of COVID-19-induced pulmonary inflammation. The purpose of this study is to evaluate the efficacy of O3FA test formulations delivered in the form of free fatty acids (FFA) or ethyl esters (EE) or as components of egg phospholipids, when administered to Wistar rats via nebulization. We also combined O3FA with common drugs used to treat pulmonary inflammation to evaluate synergy or interactions.

## MATERIALS AND METHODS

The present study was approved by the Institutional Animals Ethics Committee (IAEC) of the Palamur Biosciences Private Limited (PBPL), Proposal No. PAL/IAEC/2020/12/01/31 dated 15 May, 2021. The PBPL is a preclinical Contract Research Organization (CRO) which is certified by the Committee for the Purpose of Control and Supervision of Experiment on Animals (CPCSEA) for breeding and experimentation.

### Test ingredients and Formulation

The test O3FA took two forms: (FFA) or ethyl ester (EE). FFA were the kind gift of K.D. Pharma (Bexbach, Germany) and contained major fatty acids 31.6% EPA, 31.6% DHA, and 15.4% DPA (omega-3) and referred to below as “O3FFA”. The EE was a fish oil concentrate (Incromega E3322, Croda, United Kingdom) with 22% DHA and 33% EPA. Egg phospholipids were used as sole API and as emulsifier (Lipoid E 80, Ludwigshafen, Germany).

Montelukast Sodium (Melody Healthcare, Maharashtra, India), melatonin (Swati Spentose Pvt. Ltd., Gujarat, India), Cannabidiol (Biophore, Hyderabad, India), sodium hydrogen carbonate (Merck, India), glycerol (Merck, Germany), and Budesonide Respules™ (Pulmicort, AstraZeneca, Bangalore, India) where all obtained from the respective vendors.

Two control groups were used: G1 (Cn), a **normal control** group that received saline injections and G2 (Cd), a **disease control** group that received LPS injections similar to all experimental groups.

G3 (EPL), egg lecithin only dispersion

G4 (O3), O3FFA (50 mg/mL)

G5 (O3-0.5), O3FFA (25 mg/mL) (half dose of G4);

G6 (O3EE), O3EE (50 mg/mL) (compare to G4)

G7 (B-Ref), Budesonide Respules 0.25 mg/mL

G8 (Mont), Montelukast 4 mg/mL

G9 (MelO3), Melatonin (1 mg/mL) + O3FFA (50 mg/mL) (compare to G4)

G10 (MontO3), Montelukast (4 mg/mL) +O3FFA (50 mg/mL) (compare to G4 and G8)

G11 (CannO3), Cannabidiol (8 mg/mL) + O3FFA 50 mg/mL

Group details are provided in Table 1. Detailed compositions are presented in Supplementary Tables 1 to 6. EPL-egg phospholipids, FFA-free fatty acids, EE-ethyl esters

**Table 1.** The control, O3FA test and reference treatments grouping. After randomization, rats were divided into 11 groups (n=6). Animals from each group received the respective treatment twice daily for 7 days (3 animals/group) or 14 days (3 animals/group) through nebulization. G2 to G11 received LPS once daily. The concentration and the dose volume used for nebulization for each group are given in below table:

|  | Group | Formulation/<br>Composition | Concentration of<br>solution | Volume used for<br>Nebulization | Nebulization<br>duration |
| --- | --- | --- | --- | --- | --- |
| Cn | G1 | Control, normal; saline injection, no LPS | NA | 1.5mL | 2.5min |
| Cd | G2 | Control disease | NA | 1.5mL | 2.5min |
| EPL | G3 | Egg Lecithin only | NA | 1.5mL | 3.0min |
| O3 | G4 | O3 | 50 mg/mL | 1.5mL | 4.5min |
| O3-0.5 | G5 | O3 | 25 mg/mL | 3.0mL (1.5mL O3<br>25mg/ml + 1.5mL<br>normal saline) | 6.0min |
| O3EE | G6 | O3 ethyl esters | 50 mg/mL | 1.5mL | 3.0min |
| B-Ref | G7 | Budesonide Respules, reference therapy | 0.5 mg/mL | 1.5mL | 3.0min |
| Mont | G8 | Montelukast sodium | 4 mg/mL | 1.5mL | 3.0min |
| MelO3 | G9 | Melatonin + O3 | 1.0 mg/mL + 50<br>mg/mL | 1.5mL | 4.5min |
| MontO3 | G10 | Montelukast sodium + O3 | 4 mg/mL + 50 mg/mL | 3.0mL (1.5mL<br>montelukast sodium<br>4mg/ml+ 1.5mL O3<br>50mg/ml) | 6.5min |
| CannO3 | G11 | Cannabidiol + O3 | 8 mg/mL + 50 mg/mL | 1.5 mL | 4min |

Note: Cn- normal control; Cd- disease control; EPL- Egg phospholipids; O3- Omega-3 free fatty acids; O3EE- Omega-3 ethyl esters; B-Ref- Budesonide reference, Mont- Montelukast sodium; MelO3- Melatonin + O3; MontO3- Montelukast sodium + O3; CannO3- Cannabidiol + O3

### Animals

Male Wistar rats, a common model for acute lung inflammation [45, 46], were grouped based on a stratified randomization method and were acclimatized for five days from arrival to testing. Three animals per cage were housed in polypropylene rat cages with corn cob bedding. All the animal cages were identified by cage cards and the corresponding individual animal numbers were marked with marker pen on the base of the tail. Animals were fed standard rat chow and provided water *ad libitum*.

A total of 66 animals were used (10% extra animals were taken for randomization). Housing facilities were maintained at 19.7-22.0°C with 44 −60% relative humidity, 12-h light/dark cycle and a minimum of 12-15 room air exchanges per hour. Body weights of the animals at the time of dosing ranged from 138.9 – 169.2 g.

### Animal grouping, induction of lung inflammation and treatment

After acclimation, rats were divided into 11 groups with n=6 rats in each group. Mornings, the protocol was initiated by an intraperitoneal injection of LPS (2 mg LPS per mL prepared fresh daily, dose of 2 mg per kg body weight) for groups G2 to G11. Group G1 received a saline injection. One hour later, animals were restrained and the test item was administered by inhalation using a standard consumer nebulizer (Romsons Angel™ Nebulizer compressor system, GS-9023, Uttar Pradesh, India). Approximately 7 hours later, the animals were again restrained and treated by inhalation, and then released back to their cages for the night. Volume used for nebulization and the nebulization duration are provided in Table 1. One LPS dose and two bouts of test article inhalation, morning and evening, were repeated daily for 7 days for half (n=3) of the animals in each group and then they were euthanized by CO_2_ inhalation. The protocol was repeated for 14 days for the remaining n=3 animals, at which time they were sacrificed. Thus, animals from each group received the respective treatment (Table 1) twice daily for 7 days (3 animals/group) or 14 days (3 animals/group) through nebulization. Animals were observed for overt signs of distress during entire period of dosing and until termination of study, and also observed for prompt effects during and after administration of inhaled lipids.

Technicians performing the experiments were blinded to the treatment/reference and knew only group numbers.

On Day 8 and Day 15 three animals from each group were euthanized, lungs were harvested, weighed and bronchoalveolar lavage fluid (BALF) was collected from left lung. Harvested lungs were preserved in 10% neutral buffered formalin (NBF) and processed for histopathological evaluation. Before euthanasia blood was also collected and plasma was separated for further analysis.

#### Histopathology

Formalin fixed left lung tissue samples were processed for histopathology following standard procedures. Tissues were embedded in paraffin blocks and sections were stained routinely with Hematoxylin and Eosin staining (H&E) procedure. For each animal one slide was prepared and stained with H&E stain. Full histopathology was performed on the preserved lung tissues of all animals in the control and treatment groups. All gross lesions were examined. The lung tissue sections of 3-5 micron were stained with H&E stain and were microscopically analyzed using microscope (Make: Leica DM1000 LED; Camera: Leica MC170HD). The images were taken with 10X objective lens (Magnification X 100) using Leica LAS V4.12 software.

#### Immune Markers

The collected BALF samples were analyzed for the IL-6, IL-1β and TNF-α levels for all the animals. IL-10 and TGF-β were estimated in the plasma samples in all the animals. ELISA kits (Krishgen BioSystems, Mumbai, India) were used for the analysis of different immune markers (Suppl. Table 7).

#### Statistical analysis

Immune data are expressed as Mean ± SD. LPS disease control group (G2) was compared with all the treatment groups (G3-G11) using Student’s t-test. p<0.05 was set as statistical significance threshold.

## RESULTS

Body weights were recorded on the day of the arrival, on the day of randomization and, on the day of dosing for all groups. All animals increased body weight during the study period (Suppl. Table 8). Similarly, no differences were found in lung weights between Cd and treatment or reference groups. No mortality was observed during the acclimatization or study period (Suppl. Table 9).

Animals from groups O3EE, Melatonin+O3 and Cannibidiol+O3 showed eye irritation and lacrimation immediate after nebulization, apparently due to nebulization fumes reaching the eyes. These clinical signs subsided spontaneously within 30 minutes after nebulization. No other anomalous clinical symptoms were noted.

### Histopathology Evaluations

External and internal pathological examination of the lung tissues did not reveal any gross abnormalities in any of the control or treatment groups.

Microscopic Findings. LPS induced specific changes in the disease control group compared to the normal control, treatment and reference groups (Suppl. Figure 1).

Alveolar histiocytosis was scored as 0 (none) and by increasing severity as 1, 2, or 3. Scores were added and then normalized on a scale of 0 (normal) to 1 (maximum). All animals in the control disease (Cd) group had at least minimal macrophage infiltration leading to the highest severity score in the study of 0.61 (Figure 1). All treatments had reduced alveolar histiocytosis scores. Budesonide (B-ref) animals had the lowest score, based on one animal scored as slight and the rest normal. The treatments containing O3 scored less than half severity compared to Cd.

**Figure 1.**
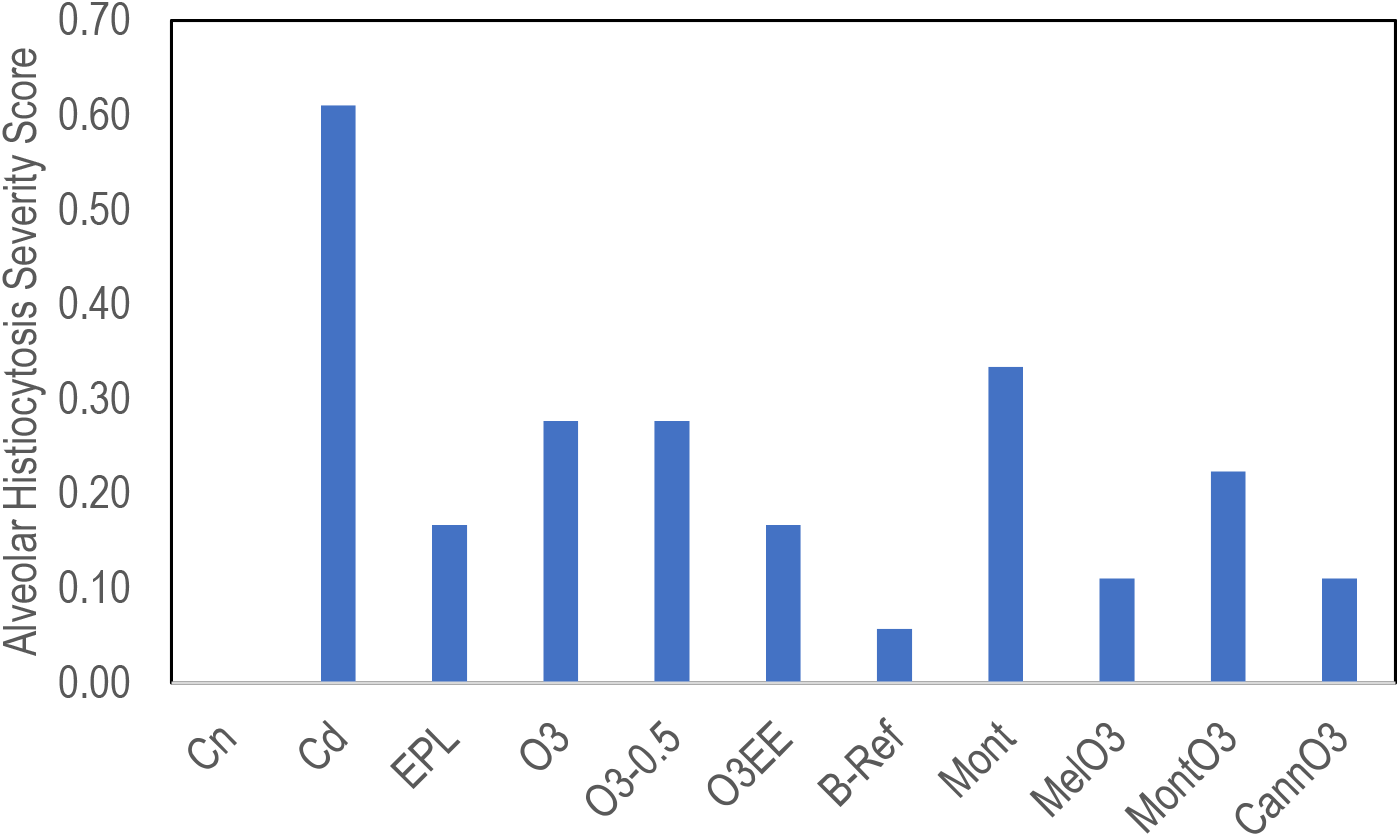
Represents alveolar histiocytosis severity scores. All animals in the control disease (Cd) group had the highest severity score (0.61). Reduced alveolar histiocytosis scores are seen in all the treatment groups. B-ref group had the lowest score compared to all treatment groups. The O3 treatment groups scored less than half severity compared to Cd.

Minimal vacuolation was observed in 1-2 animals in the EPL, O3, O3-0.5, and O3EE groups.

#### Immune Parameters

BALF samples were analyzed for the IL-6, IL-1β and TNF-α levels for all the animals. IL-10 and TGF-β were estimated in the plasma samples of all the animals.

The 8 day and 15 day means for each group were tested for significant differences by pairwise t test. IL-6, TNFα, and TGF-β were not significantly different for time points in any of the groups, thus results were pooled as n=6. Several groups were significantly different for IL-10 and IL-1β, and were analyzed at each time point with n=3. Comparisons between groups were done within each time point.

Results of IL-6, TNF-α and TGF-β are presented in Figure 2 as mean ± SD.

**Figure 2.**
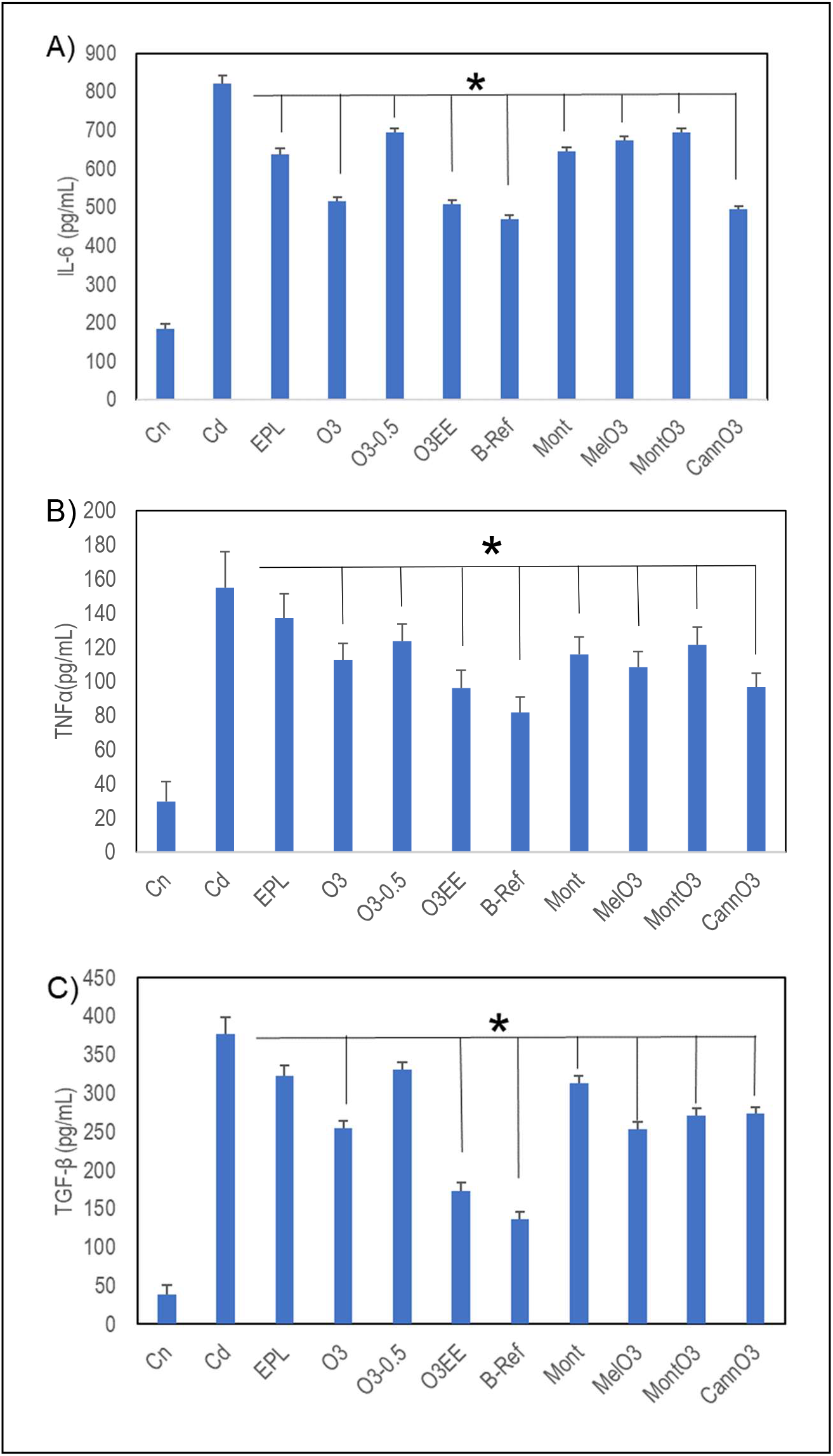
Cytokine levels in treatments compared to the disease control Cd (mean±SD). No time effects were found for these groups; day 8 and day 15 results were pooled to yield n=6. A) IL-6 levels are downregulated and reached statistical significance in all the treatment groups and B-Ref by pairwise comparison (*p<0.05). B) TNF-α levels are downregulated and reached statistical significance in all the treatment groups and B-Ref (*p<0.05), except EPL. C) TGF-β levels are downregulated; all significantly different from Cd (*p<0.05), except EPL and O3-0.5 groups.

When compared to disease control group (Cd), IL-6 levels are significantly downregulated in all the treatment groups and reference B-Ref (Figure 2A). O3, O3EE, and CannO3 had lowest IL-6, similar to the Budesonide Respules reference (B-Ref). TNF-α is significantly downregulated in all the groups except EPL (Figure 2B). The O3EE and CannO3 were similar to Budesonide reference (B-Ref). TGF-β is significantly downregulated in all groups except EPL and O3-0.5 (Figure 2C). O3EE attenuation of TGF-β was similar to the reference Budesonide (B-Ref).

Most of the O3 groups were significantly lower in IL-10 levels than Cd at the two time points (Figure 3A). O3-0.5, CannO3, and B-Ref increased with time, whereas O3EE decreased. By pairwise comparison with Cd at identical time points, O3, O3EE, B-Ref, Mont, MelO3 and CannO3 are significantly lower at both time points; compared to Cd, O3-0.5 is significantly different only at day 8 and MontO3 is different only at day 15. EPL was not significant at either time point. In contrast, few of the treatments were significantly different for IL-1β (Figure 3B). O3EE was the only treatment with lower IL-1β than Cd at both time points. B-Ref, MelO3, and CannO3 were significantly lower only at 15 days.

**Figure 3.**
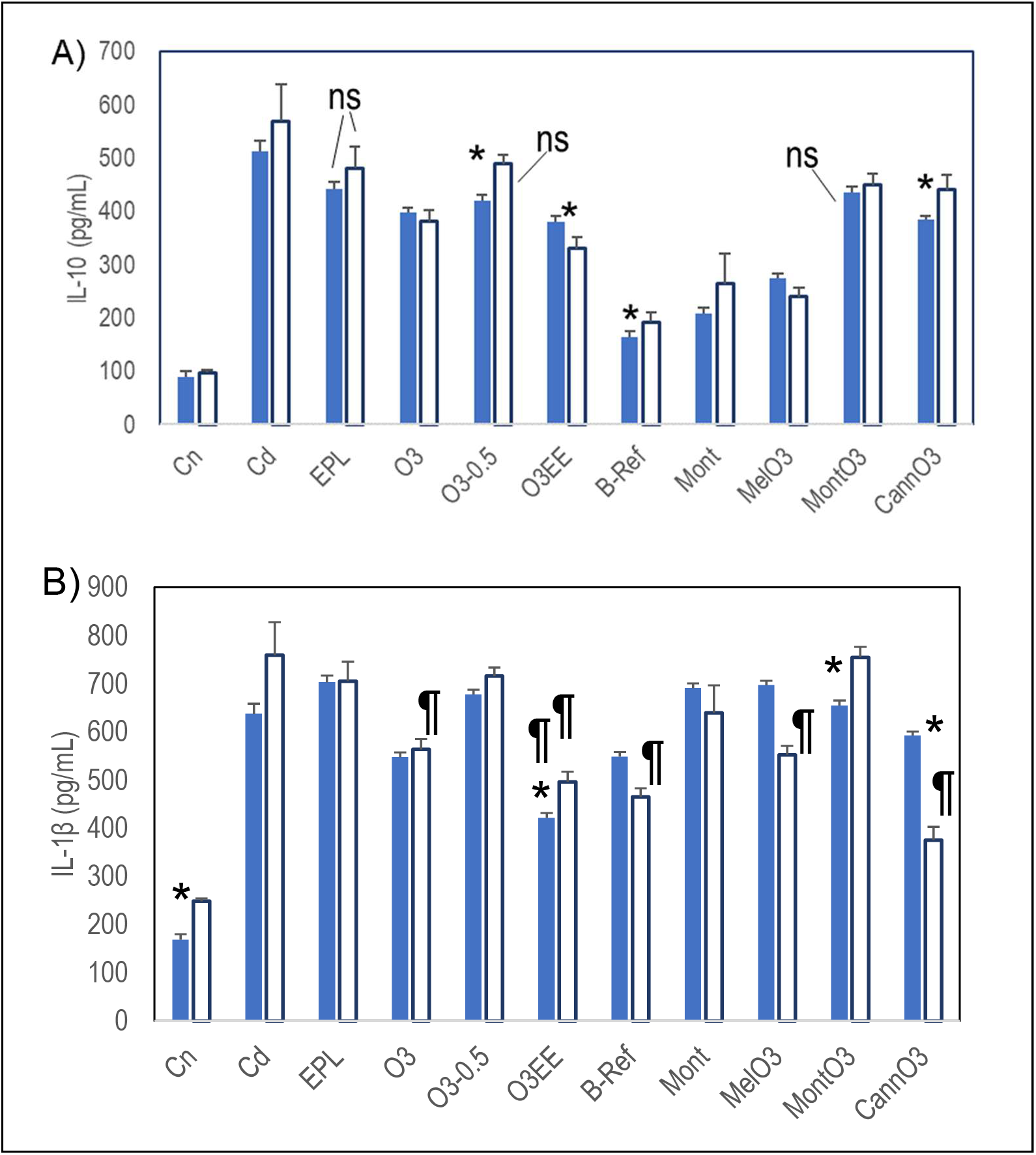
Cytokine levels in treatments compared to the disease control Cd (mean±SD). Time effects were significant for these groups; day 8 results were compared to day 8, and day 15 to day 15, n=3 per group. A) With time IL-10 for O3-0.5, CannO3, and B-ref increased, whereas O3EE decreased (*p<0.05 for time effect). All comparisons at single time points are significant except EPL at both time points, O3-0.5 at day 15 and MontO3 at day 8 (“ns”). B) IL-1β levels were mostly not significantly different from Cd except those labeled with ¶. O3EE is the only treatment significant at both the time points. O3, B-Ref, MelO3 and CannO3 are significant only at the 15 day time point.

## DISCUSSION

We demonstrate here for the first time O3FA formulations delivered via nebulization reduce LPS-induced acute lung inflammation. Histopathology microscopic observations and immunological marker analysis revealed significant reduction of inflammation in O3FA treatment groups compared to disease control (Cd) group.

Our O3FA were delivered primarily in the form of FFA to enhance incorporation into lung tissue and minimize lipoid pneumonia. The single treatment group using ethyl esters (O3EE) was in the most common form of oral O3 supplements. No symptoms related to excess lipid accumulation was observed, and the O3EE treatment was among the most effective in treating effects of LPS.

We are aware of only one other study of an inhalation form of any O3FA. Inhaled ovalbumin was used to model atopic asthma in mice. Aerosolized tridocosahexaenoyl-glycerol (DHA-TG) inhaled immediately prior to the ovalbumin insult significantly reduced the total eosinophil percentage in lavage fluid. The TAG-DHA did not cause lipoid pneumonia [47]. Considering the efficacy of O3EE and absence of adverse symptoms, we conclude that O3EE are a viable delivery form of nebulized O3.

Alveolar histiocytosis is characterized by the presence of foci of alveolar macrophages and mixtures of inflammatory cells [48]. Histopathology microscopic observations revealed minimal to moderate alveolar histiocytosis in all 6 LPS-treated disease control (Cd) animals (Suppl. Figure 1). None of the animals in the treatment or reference groups had moderate alveolar histiocytosis, however, minimal to slight alveolar histiocytosis was noted in the O3FA treatment and reference groups (Suppl. Figure 1). When compared to the O3 FFA treatment groups (O3 and O3-0.5), O3EE, B-ref, MelO3 and CannO3 groups showed better alveolar histiocytosis and inflammation resolution. In the Cd disease control and Mont groups 3 out of 6 animals showed slight alveolar histiocytosis. Thus, montelukast had less impact on the alveolar histiocytosis and inflammation resolution. Multifocal alveolar histiocytosis was seen in O3 and O3-0.5 group animals.

The pro-inflammatory synovial phospholipase A2 (PLA2) was found to be highly elevated during acute lung injury and in LPS stimulated guinea pig alveolar macrophages [49–51]. High amounts of pro-inflammatory omega-6 arachidonic acid (AA) suppress anti-inflammatory omega-3 DHA, DPA and EPA synthesis and accumulation in cell membranes. The AA derived eicosanoids regulate immunopathological processes ranging from inflammatory responses to tissue remodeling [52, 53]. AA-derived prostaglandin (PG) synthesis occurs upon release of AA within the membranes by PLA2 [54], thus PG synthesis is limited by the supply of AA. DHA that substitutes for AA is a known inhibitor of PG synthesis by cyclooxygenase [55]. Dietary EPA and AA compete for incorporation into membrane phospholipids but also for biosynthesis from their respective FA precursors, as they share same enzyme for their biosynthesis [56, 57]. AA derived metabolites mediate inflammation in lung disorders [11, 58, 59], whereas, DHA, DPA and EPA derived metabolites resolve inflammation and clotting [11, 60–64]. Alveolar histiocytosis was present in 100% of SARS-CoV-2 infected cats [65]. Resolution of lung inflammation and thrombosis is the main purpose of our new O3FA formulations.

COPD caused 3.23 million deaths in 2019 and is the third leading cause of death worldwide [66]. Elevated IL-6 levels in the exhaled breath condensate samples are associated with airway inflammation in COPD patients [67]. TNFα over-expression in both humans and animal models showed pathological changes consistent with both emphysema and pulmonary fibrosis. Mice lung histology and computed tomography images showed changes involving airspace enlargement, loss of small airspaces, increased collagen and thickened pleural septa [68]. Increased expression of TGF-β is seen in lung specimens collected from COPD patients [69]. IL-10 levels are elevated in COPD patients [70]. Elevated serum IL-1β levels are associated with airway inflammation in COPD patients [71]. Higher intake of omega-3 fatty acids is associated with lower risk of severe exacerbations, better health-related quality of life, and fewer respiratory symptoms in COPD patients [37].

Several studies have shown IL-6 as the critical tumor-promoting cytokines in NSCLC. IL-6 levels are increased in the serum and exhaled breath condensate samples from NSCLC patients and are related to tumor size [72]. By inducing epithelial-mesenchymal transition of lung cancer cells IL-6 and TNF-α can promote invasion and metastasis in NSCLC [73]. Increased TGF-β expression was found to be associated with lymph node metastasis and tumor angiogenesis in NSCLC [74]. In late stage NSCLC patients increased expression of IL-10 is seen in tumor-associated macrophages [75]. IL-1β is a key mediator of the initiation of inflammatory response in NSCLC and a potent inducer of the COX2–PGE2 pathway, leading to immune suppression [76]. Cachexia is frequently observed in lung cancer; omega-3 oral supplementation preserved body weight in NSCLC patients undergoing chemoradiotherapy [38]. Lung cancer patients whose plasma phospholipid EPA concentrations were higher showed better preservation of body weight [38]. Most of our O3FA test treatments reduced the levels of IL-6, TNF-α, TGF-β, IL-10 and IL-1β significantly (Figures 2 and 3).

Multiple studies have shown elevation of both pro-inflammatory and anti-inflammatory cytokines in COVID-19 patients, reviewed by Dhar et al [77]. Dysregulation and increased production of IL-1β and its downstream molecule IL-6 are seen in severe COVID-19 patients [78, 79]. IL-6 and IL-10 are found to be predictive of COVID-19 disease severity [80]. A dramatic elevation of IL-6 and IL-10 levels is a characteristic feature of the cytokine storm in COVID-19 patients [81–83]. Persistent viral stimulation, and IL-6, IL-10 and TNF-alpha levels, are indicators of T-cell exhaustion in COVID-19 patients [84]. Increased IL-6 and TNF-α levels are significant predictors of COVID-19 severity and death [85].

In COVID-19 patients, increased pro-inflammatory IL-6 levels are associated with increased body temperature, elevation in CRP and ferritin inflammation markers, pulmonary inflammation and extensive lung damage [77, 86]. IL-6 levels are elevated in COPD patients [67], asthma patients [87, 88] and other inflammatory lung disorders [88]. All our O3FA test treatments reduced the levels of IL-6 significantly (Figure 2A). Omega-3 HUFA reduce plasma concentrations of IL-6 and ameliorate systematic inflammation [89]. Ma et al. 2016 found that an allele of IL6 rs2961298 SNP was associated with higher cg01770232 methylation and increased IL-6 levels, however, higher circulating omega-3 HUFA concentration by interacting with rs2961298 reduced cg01770232 methylation and IL-6 levels [90]. Intratracheal DHA pre-treatment reduced IL-6 levels and mitigated bleomycin-induced pulmonary inflammation and fibrosis in a mice model [91].

Tissue necrosis factor-α (TNF-α) is upregulated in most inflammatory conditions and contributes to changes in the blood coagulation [92]. Increased TNF-α along with IL-6 and IL-10 are hallmarks of a hyperinflammatory response and an underlying cytokine storm in COVID-19 patients [93]. An excessive amount of ferritin in COVID-19 patients is also reflective of a surplus of TNF-α levels [94]. TNF-α is upregulated in several inflammatory lung disorders [68]. Fish oil treatment decreased TNF-α production in healthy human volunteers [13]. All our O3FA test treatments reduced the levels of TNF-α significantly at both the time points (Figure 2B). Notably, the O3EE and CannO3 groups were comparable to the B-Ref (Budesonide) steroid reference group.

Transforming growth factor β (TGF-β) is a pleiotropic cytokine which plays a major role in inflammatory conditions [95]. TGF-β along with IL-6 drives the differentiation of T helper 17 (Th17) cells, which promote inflammation and augment autoimmune conditions [95, 96]. In addition, TGF-β promotes the differentiation of IL-10 producing T cells, which lack suppressive function and in turn promote tissue inflammation [97, 98]. Increased expression of TGF-β cause and promote lung fibrosis in COVID-19 patients [99, 100]. Increased expression of TGF-β is seen in acute lung injury and several chronic lung inflammatory disorders [101, 102]. All our O3FA test treatments, except O3-0.5 reduced the levels of TGF-β significantly and the reduction was comparable to the B-Ref in O3EE (Figure 2C).

IL-10 is a pleiotropic cytokine whose primary function in most tissues is to limit the inflammatory response, however, in COVID-19 it is dramatically elevated. This phenomenon in COVID-19 is thought to be a negative feedback mechanism to suppress inflammation [81]. IL-10 is also known to introduce T-cell anergy during viral infection [103]. Our O3FA test treatments reduced the levels of IL-10 significantly (Figure 3A).

IL-1β is a pro-inflammatory cytokine that is crucial for host-defense responses to infection, antimicrobial immunity and autoimmune inflammation [104, 105]. IL-1β levels are associated with cytokine storm in a subset of COVID-19 patients [106, 107]. IL-1β expression levels are found to be significantly increased in the bronchial wall of asthmatic patients [108]. On the other hand, fish oil treatment decreased IL-1β production in healthy human volunteers [13]. Our O3FA test treatment O3EE reduced the levels of IL-1β significantly at both time points and the reduction was better than the B-Ref group (Figure 3B).

Multiple animal and human studies indicate that both inhaled and systemic corticosteroids cause immunosuppression and impair induction of anti-viral type-I interferon responses to a range of respiratory viruses, including COVID-19 [109–114]. These are usually undesirable side effects. In India and other regions of the world, a significant increase in the incidence of fungal infections such as invasive aspergillosis or mucormycosis, a life-threatening angioinvasive maxillofacial fungal infection(s) due to corticosteroid administration, has been reported in many individuals suffering from COVID-19, especially in patients with diabetes [114–118]. In patients with overwhelming viral illness, broad immunosuppression may be inadvisable [109]. The novel O3FA formulations thus represent an alternative therapy with some significant advantages over corticosteroids. O3FA are dietary, natural, endogenous metabolites that promote balanced inflammatory and thrombotic responses. The safety and efficacy of multigram oral and intravenous lipid emulsions has long been established in young children and adults [119–123]. A double-blind, randomized clinical trial showed oral O3FA improved the levels of several respiratory and renal function parameters in critically ill COVID-19 patients [124]. The present formulations include administration of O3FA without substantial adverse reactions or side effects.

Persons who have survived the acute disease and have long term symptoms are known as “COVID Long Haulers”. Prominent symptoms in these patients are fibromyalgia, fatigue, and sleep disturbance [125–127]. The O3FA, due to their anti-inflammatory effects, are known to be beneficial in the treatment of arthritis and neuropathic pain associated with fibromyalgia syndrome (FMS) [128–130]. Melatonin has helped in reducing anxiety, lung fibrosis and controlling insomnia in COVID-19 patients [131–135]. On this basis, our MelO3 treatment group may reduce long haul COVID-19 symptoms, in addition to lung fibrosis.

The O3FA are precursors for the synthesis of endocannabinoids, including endocannabinoid epoxides with powerful anti-inflammatory properties [136, 137]. Cannabidiol exerts a wide range of anti-inflammation and immunomodulation effects and can mitigate the uncontrolled cytokine storm during acute lung injury [138]. Nguyen et al. 2021 showed CBD administration is associated with decreased risk of SARS-CoV-2 infection in humans and can block SARS-CoV-2 infection at early stages [139]. A combined inhalation cannabidiol-O3FA modeled by our CannO3 group is thus likely to reduce runaway inflammation and thrombosis by mitigating the uncontrolled cytokine storm in COVID-19.

## CONCLUSIONS

In the present study we showed novel forms of pharmaceutical grade O3FA, specifically docosahexaenoic acid (DHA), docosapentaenoic acid (DPA) and/or eicosapentaenoic acid (EPA) medicament delivery to the rat lungs via nebulization reduced LPS induced acute lung inflammation. Both histopathology observations and immunological marker analysis revealed significant reduction of inflammation in our O3FA treatment groups compared to the disease control (Cd). COPD is a global epidemic, killing over 3.2 million individuals each year and there is no cure for it. COVID-19 is an enigma and treatment protocols are still evolving to treat critically ill patients with COVID-19. One of the major challenges during COVID-19 pandemic is to prevent disease progression from symptomatic to ICU. We propose that our natural, dietary, anti-inflammatory and pro-resolving O3FA formulations can help in preventing eicosanoid storm which can be followed by the cytokine storm. Our formulations can be used to treat inflammation and thrombosis related lung disorders, for example, asthma, COPD, cystic fibrosis, interstitial fibrosis, bronchiolitis, NSCLC and conditions related to acute respiratory distress like COVID-19.

## ACKNOWLEDGEMENTS

This work was funded and supported by Leiutis Pharmaceuticals, Hyderabad, India. All the animal phase and experiments are conducted at a full-fledged pre-clinical CRO Palamur Biosciences Private Limited (PBPL), Hyderabad, India. We thank the staff and researchers at PBPL for conducting this study.

## AUTHOR CONTRIBUTIONS

C.K., K.S.D.K., J.T.B. formulated the research questions and designed the study; N.B., A.N., S.A. conducted lab work; C.K., K.S.D.K., J.T.B. analyzed and interpreted the data; and C.K., K.S.D.K. and J.T.B. wrote the first draft and all authors approved the final draft.

## Supplementary Figures

**Supplementary Figure 1.**
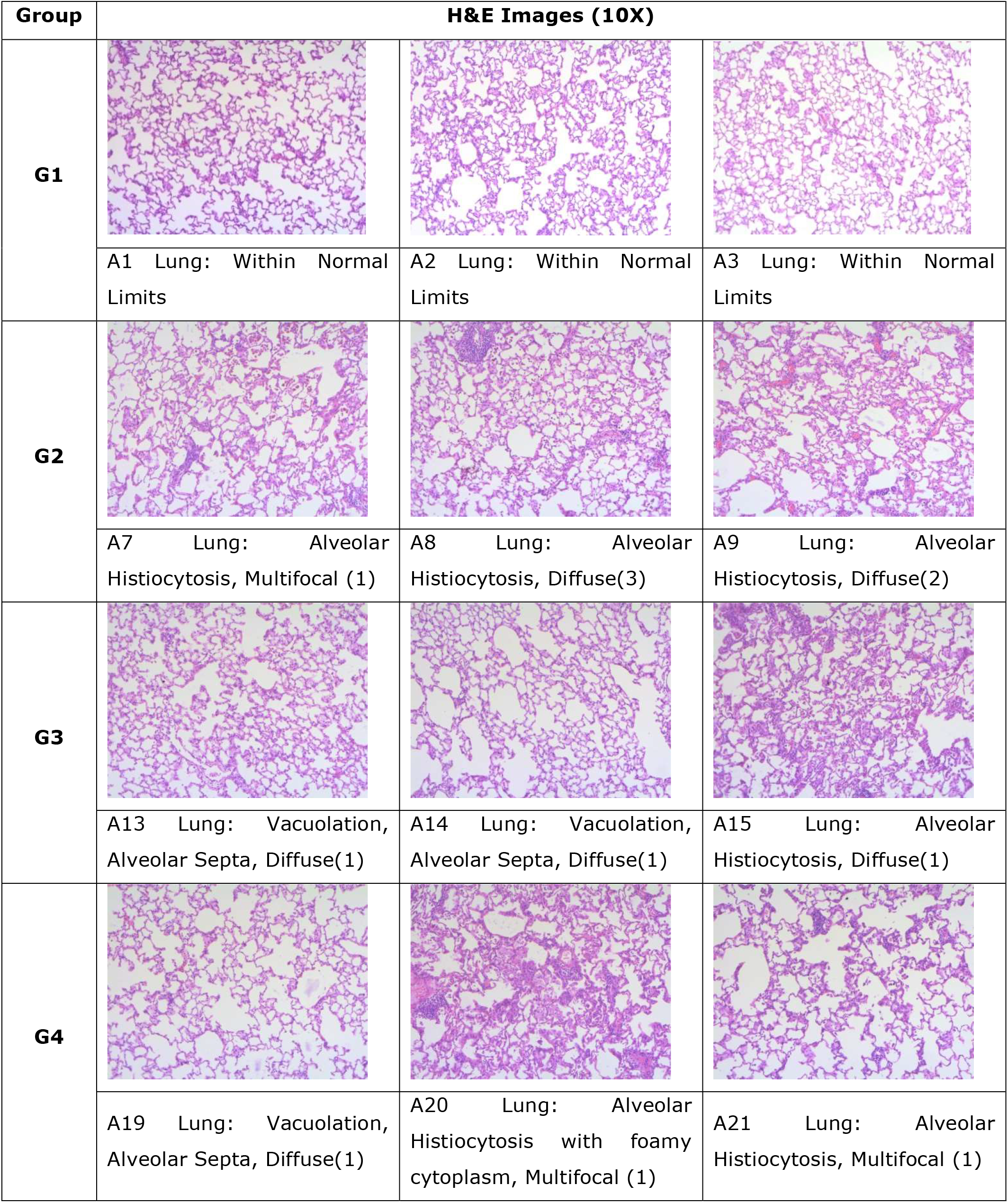
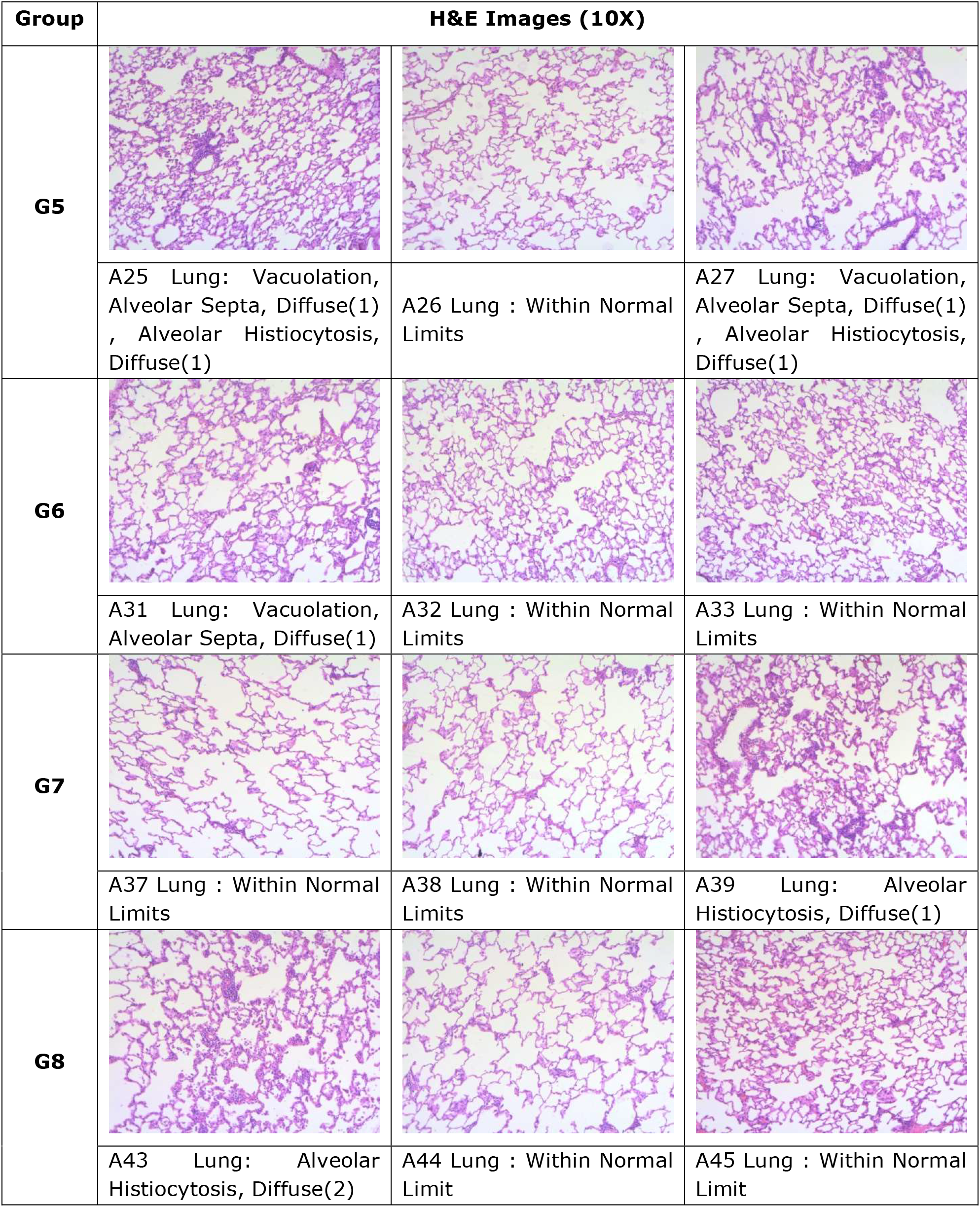
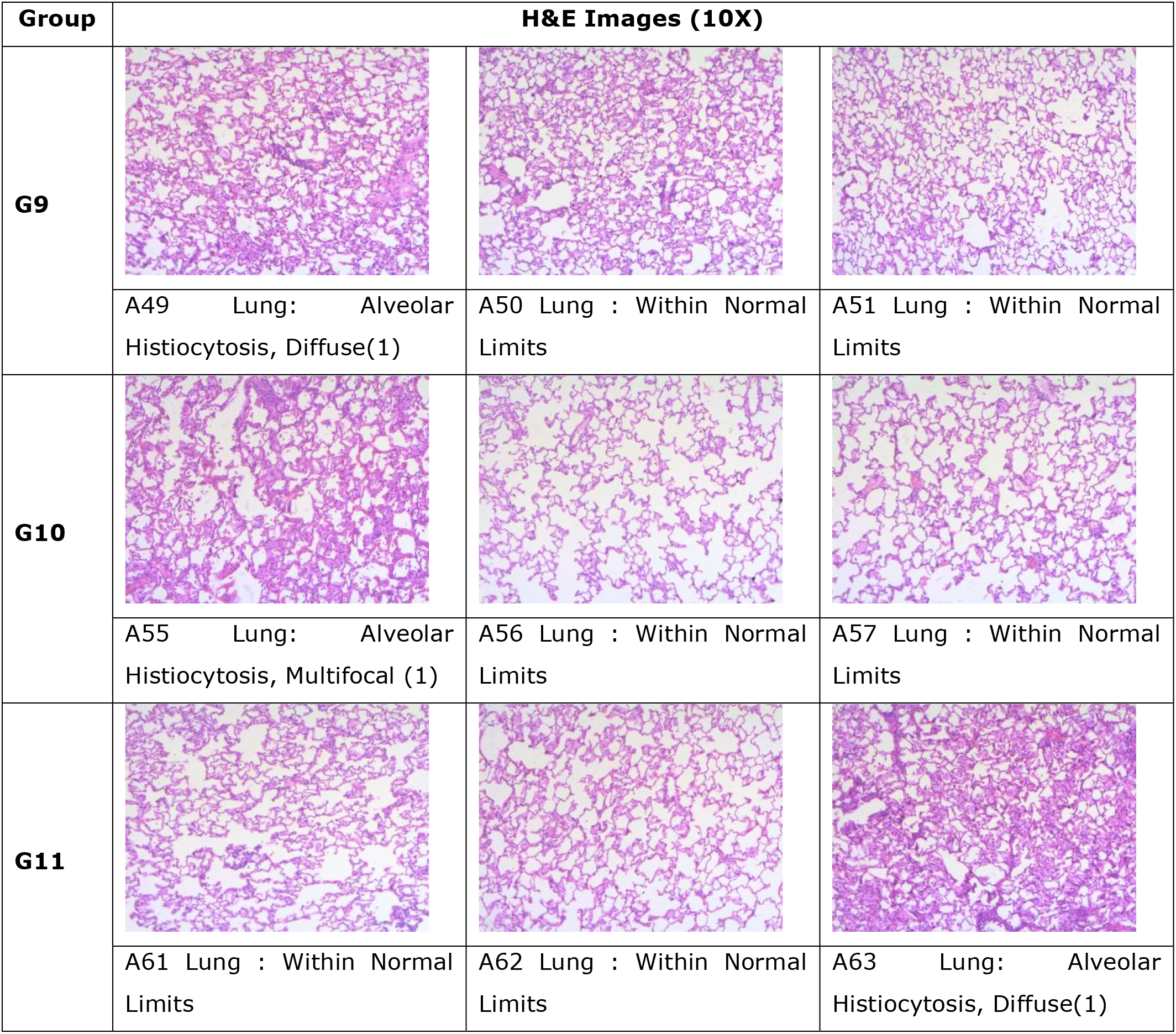
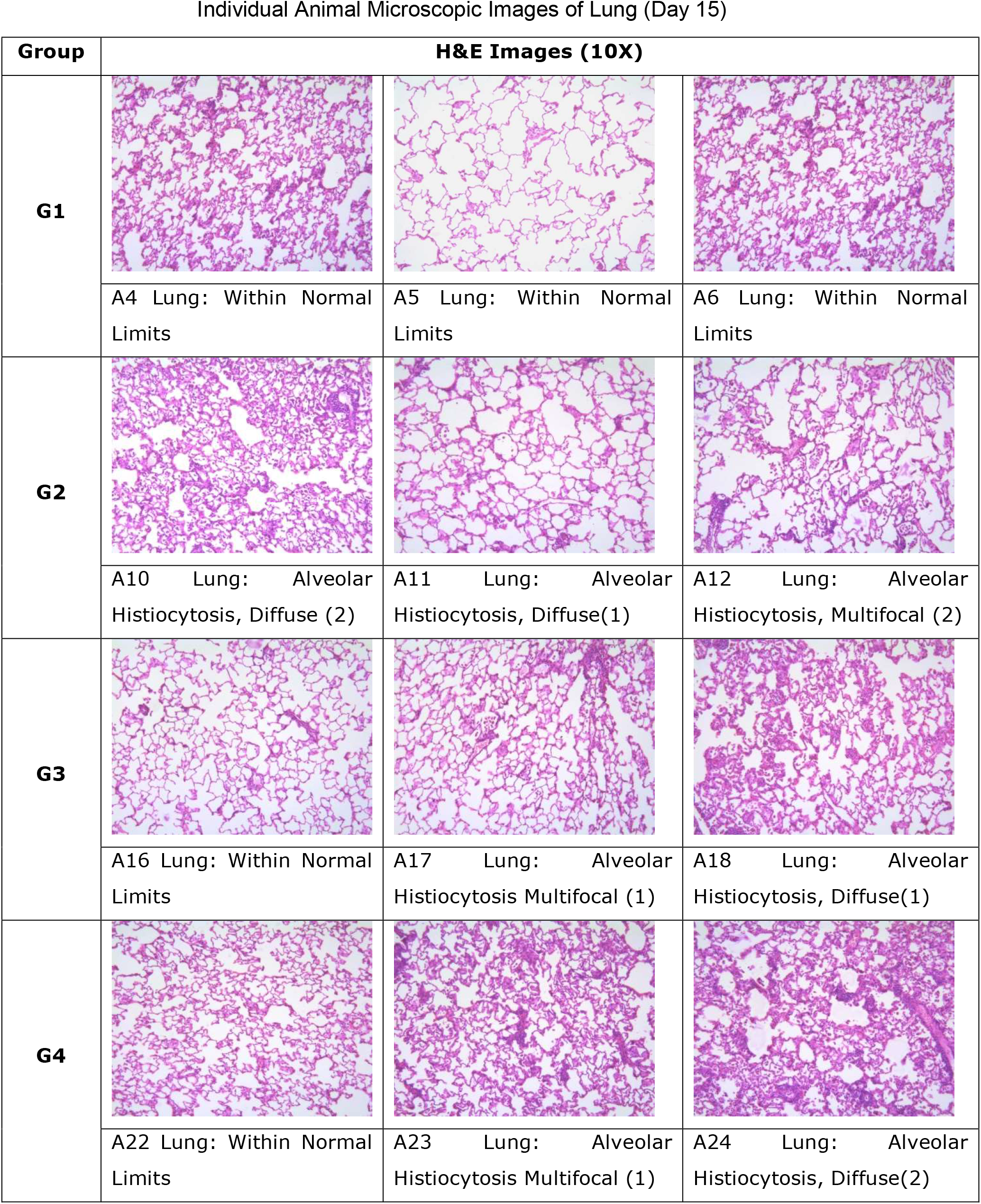
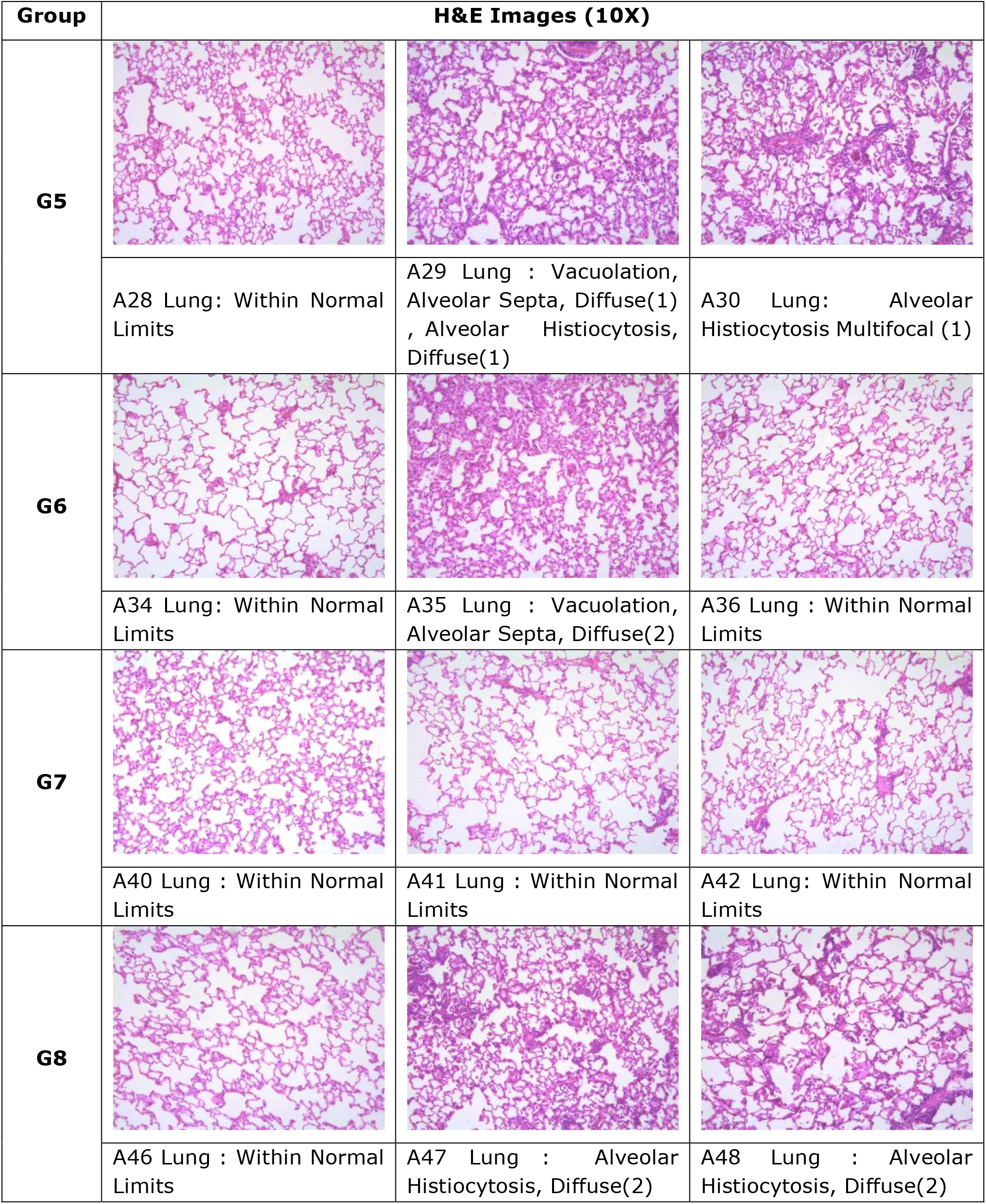
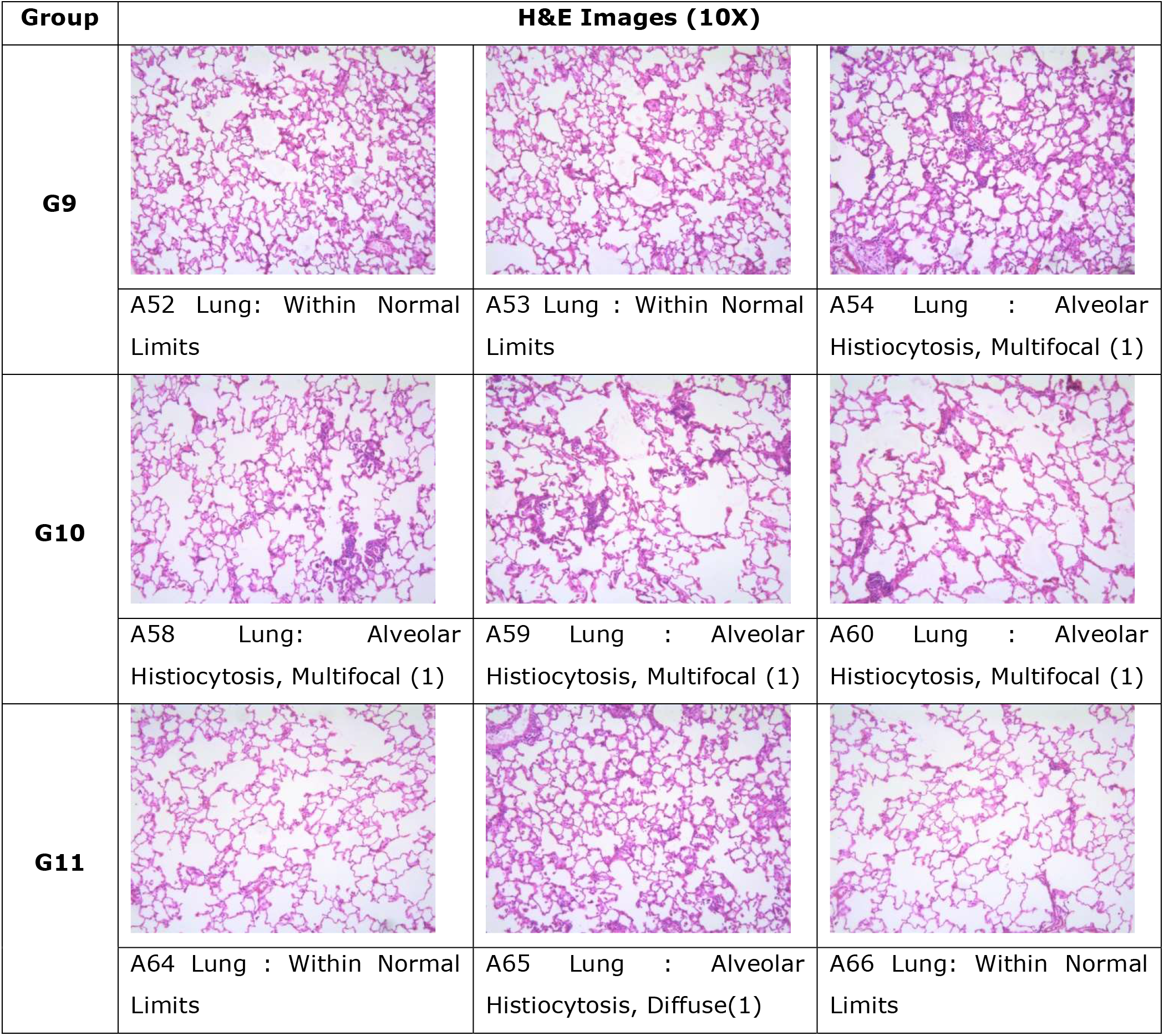
Individual Animal Microscopic Images of Lung (Day 8)

## Supplementary Tables

**Suppl. Table 1:**
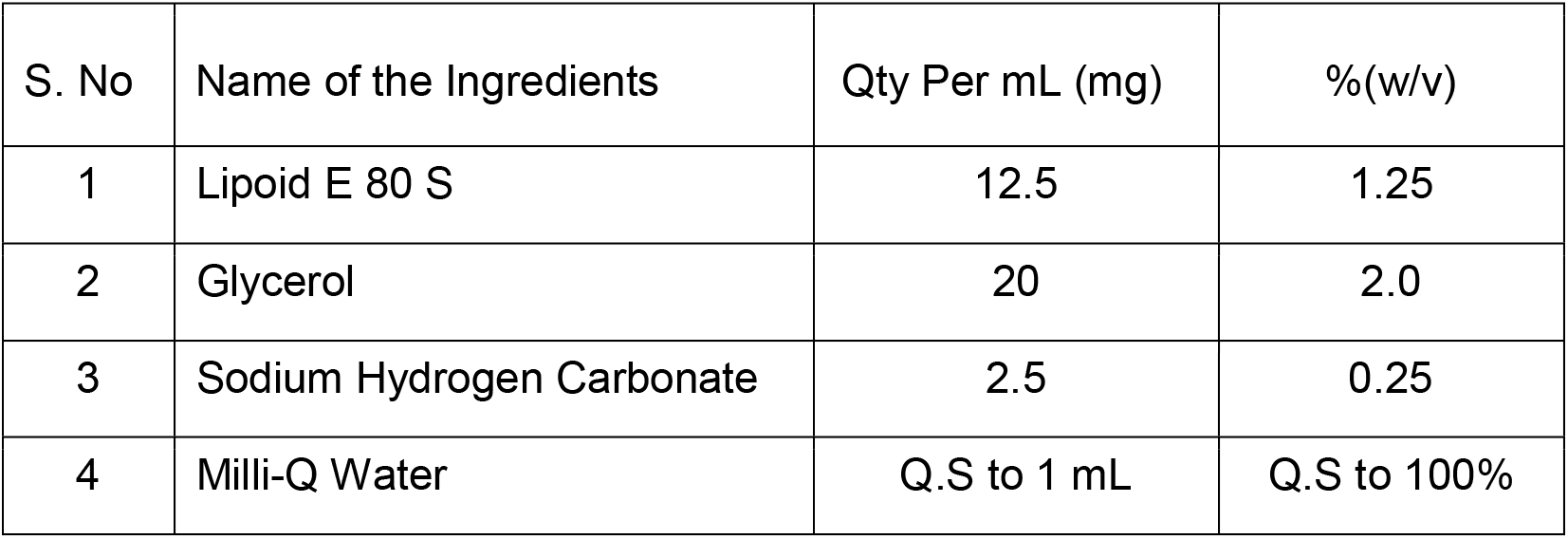
Composition of Egg Lecithin (Placebo) Emulsion (EPL group)

| S. No | Name of the Ingredients | Qty Per mL (mg) | %(w/v) |
| --- | --- | --- | --- |
| 1 | Lipoid E 80 S | 12.5 | 1.25 |
| 2 | Glycerol | 20 | 2.0 |
| 3 | Sodium Hydrogen Carbonate | 2.5 | 0.25 |
| 4 | Milli-Q Water | Q.S to 1 mL | Q.S to 100% |

**Suppl. Table 2:** Composition of Omega-3 FFA Emulsion (O3 group)

| S. No | Name of the Ingredients | Qty Per mL (mg) | %(w/v) |
| --- | --- | --- | --- |
| 1 | Omega-3 Fatty Acids | 50 | 5.0 |
| 2 | Lipoid E 80 S | 8 | 0.8 |
| 3 | Glycerol | 20 | 2.0 |
| 4 | Sodium Hydrogen Carbonate | 2.5 | 0.25 |
| 5 | Milli-Q Water | Q.S to 1 mL | Q.S to 100% |

Note: FFA- free fatty acids

**Suppl. Table 3:** Composition of Omega-3 EE Emulsion (O3EE group)

| S. No | Name of the Ingredients | Qty Per mL (mg) | %(w/v) |
| --- | --- | --- | --- |
| 1 | Fish Oil (Incromega E3322) | 50 | 5 |
| 2 | Lipoid E 80 S | 12.5 | 1.25 |
| 3 | Glycerol | 20 | 2 |
| 4 | Sodium Hydrogen Carbonate | 0.15 | 0.015 |
| 5 | Milli-Q Water | Q.S to 1 mL | Q.S to 100% |

Note: EE- ethyl esters

**Suppl. Table 4:** Composition of Montelukast Sodium Solution (Mont group)

| S.<br>No | Name of the Ingredients | Qty Per mL (mg) | %(w/v) |
| --- | --- | --- | --- |
| 1 | Montelukast Sodium | 4 | 0.4 |
| 2 | Milli-Q Water | Q.S to 1 mL | Q.S to 100% |

**Suppl. Table 5:** Composition of Melatonin Solution (MelO3 group)

| S. No | Name of the Ingredients | Qty Per mL (mg) | %(w/v) |
| --- | --- | --- | --- |
| 1 | Melatonin | 1 | 0.1 |
| 2 | Omega-3 FFA | Q.S to 1 mL | Q.S to 100% |

Note: FFA- free fatty acids

**Suppl. Table 6:**
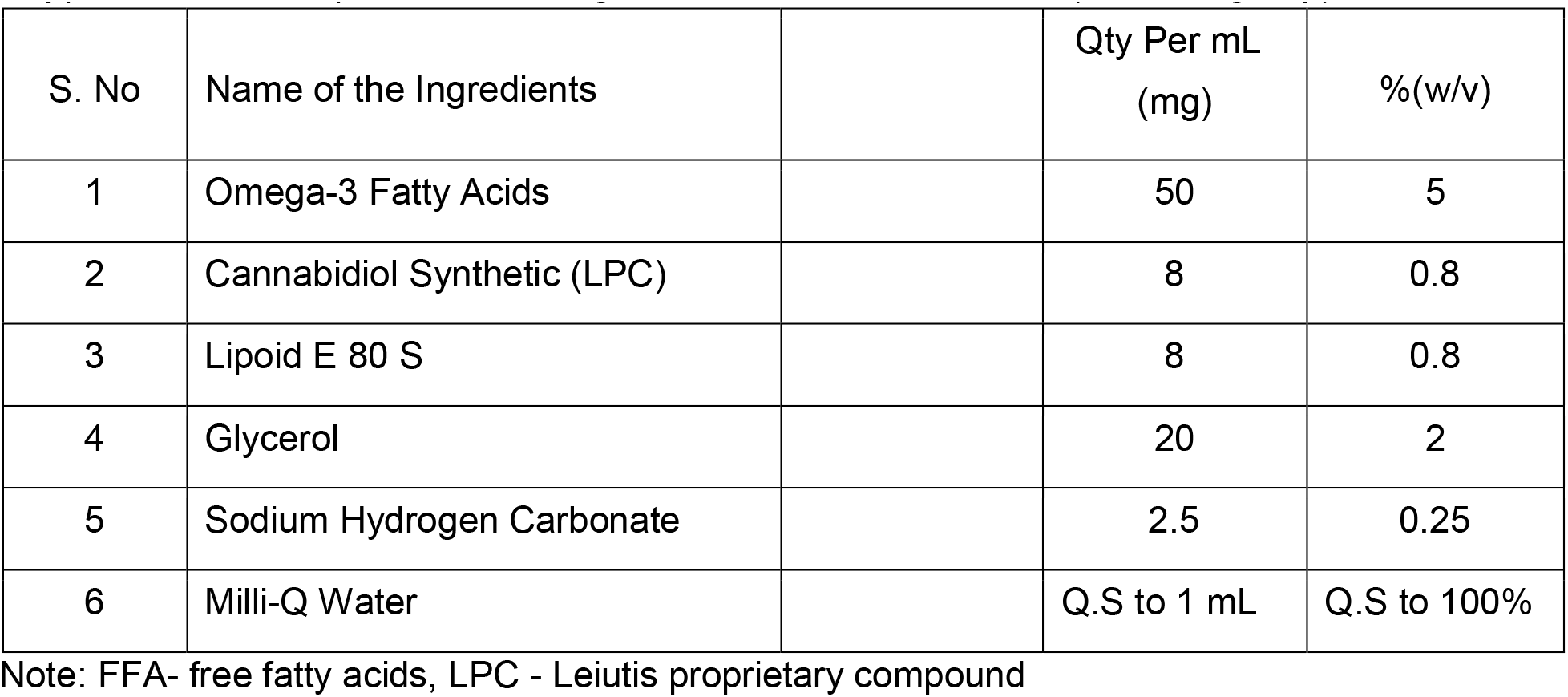
Composition of Omega-3 FFA and LPC Emulsion (CannO3 group)

| S. No | Name of the Ingredients |  | Qty Per mL<br>(mg) | %(w/v) |
| --- | --- | --- | --- | --- |
| 1 | Omega-3 Fatty Acids |  | 50 | 5 |
| 2 | Cannabidiol Synthetic (LPC) |  | 8 | 0.8 |
| 3 | Lipoid E 80 S |  | 8 | 0.8 |
| 4 | Glycerol |  | 20 | 2 |
| 5 | Sodium Hydrogen Carbonate |  | 2.5 | 0.25 |
| 6 | Milli-Q Water |  | Q.S to 1 mL | Q.S to 100% |

Note: FFA- free fatty acids, LPC - Leiutis proprietary compound

**Suppl. Table 7.** ELISA kits used to measure immune markers.

| S. No. | ELISA Kit Name | REF ID | Lot No. | Company Name |
| --- | --- | --- | --- | --- |
| 1 | GENLISA™ Rat TGF-β ELISA | KLR0778 | RTGFB0621 | Krishgen BioSystems, Mumbai |
| 2 | GENLISA™ Rat IL-6 ELISA | KB3068 | RI60621 |  |
| 3 | GENLISA™ Rat IL-10 ELISA | KLR0108 | RIL100621 |  |
| 4 | GENLISA™ Rat IL-1β ELISA | KB3063 | RTB0721 |  |
| 5 | GENLISA™ Rat TNF-α ELISA | KB3145 | RTA0521 |  |

**Suppl. Table 8.**
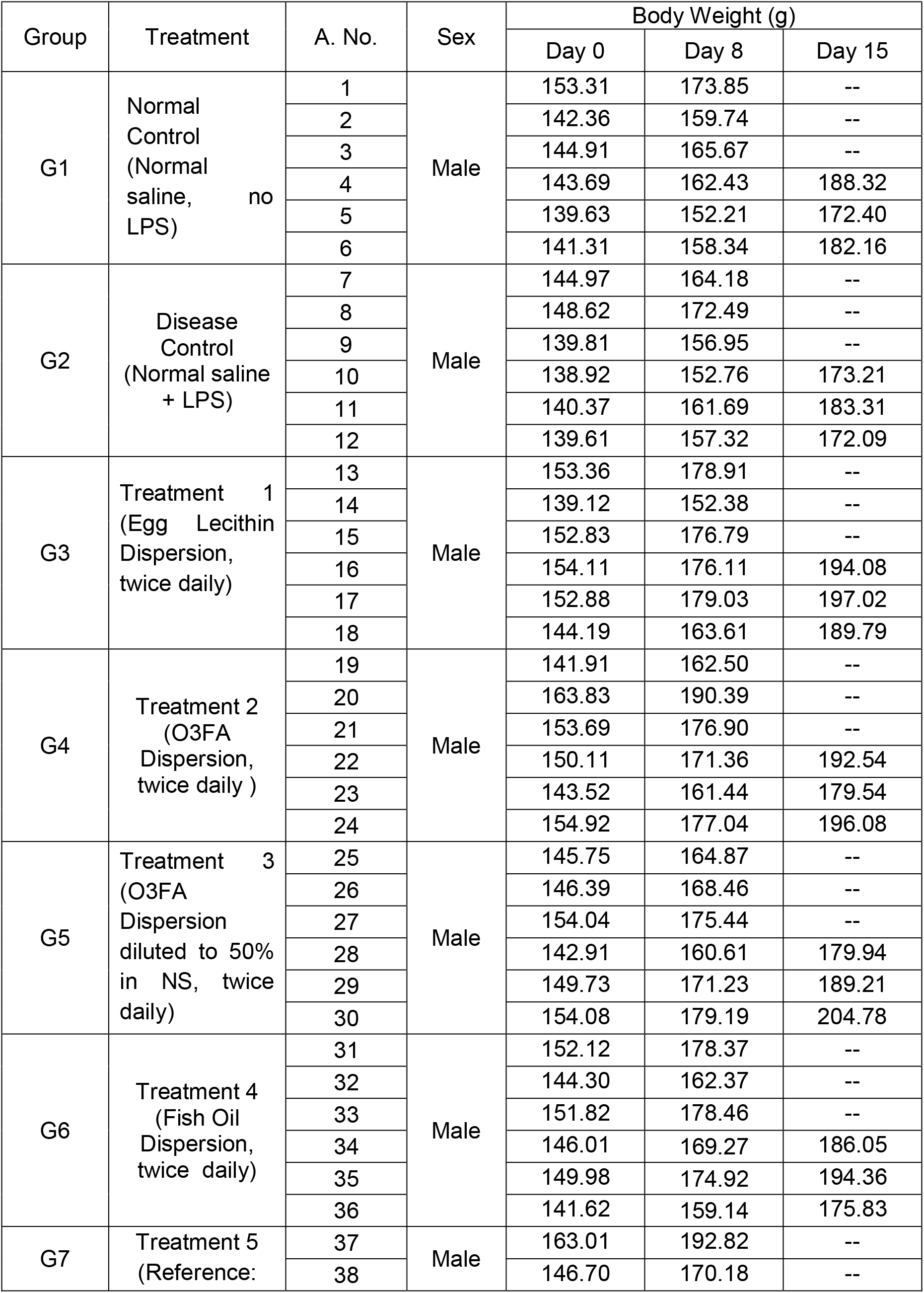
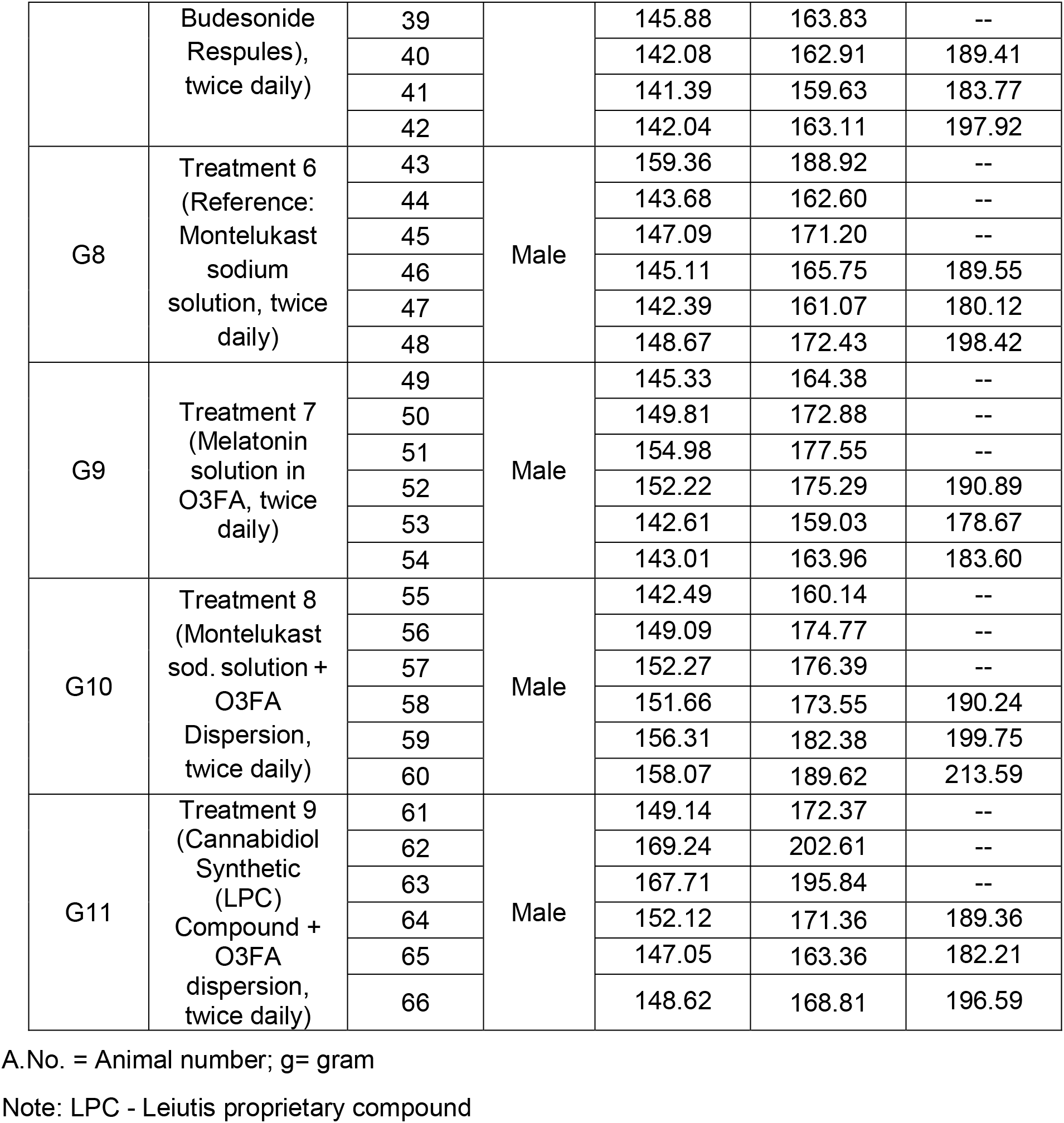
Individual Animal Body Weights.

**Suppl. Table 9.** Summary of Mortality.

| **Observation Period** | **Parameter** | **Sex** | **Animal Numbers** | **Incidences** |
| --- | --- | --- | --- | --- |
| Acclimatization Phase Day (1-4) | Mortality | Male | 1 -72 | 0/72 |
| Treatment / Experiment Phase (Day 0-7) | Mortality | Male | 1-66 | 0/66 |
| Treatment / Experiment Phase (Day 8-14) | Mortality | Male | 33 | 0/33 |

0/72 – 0 Mortality out of 72 animals

## References

[1] S. Han, R.K. Mallampalli, The acute respiratory distress syndrome: from mechanism to translation, J Immunol, 194 (2015) 855–860.

[2] S.F. Pedersen, Y.C. Ho, SARS-CoV-2: a storm is raging, J Clin Invest, 130 (2020) 2202–2205.

[3] D. Claar, T.V. Hartert, R.S. Peebles, Jr., The role of prostaglandins in allergic lung inflammation and asthma, Expert Rev Respir Med, 9 (2015) 55–72.

[4] R.J. Martin, Therapeutic significance of distal airway inflammation in asthma, J Allergy Clin Immunol, 109 (2002) S447–460.

[5] E.A. Roesch, D.P. Nichols, J.F. Chmiel, Inflammation in cystic fibrosis: An update, Pediatr Pulmonol, 53 (2018) S30–S50.

[6] P.J. Barnes, Inflammatory mechanisms in patients with chronic obstructive pulmonary disease, J Allergy Clin Immunol, 138 (2016) 16–27.

[7] S.J. Bourke, Interstitial lung disease: progress and problems, Postgrad Med J, 82 (2006) 494–499.

[8] B.L. Yen, M.L. Yen, L.T. Wang, K.J. Liu, H.K. Sytwu, Current status of mesenchymal stem cell therapy for immune/inflammatory lung disorders: Gleaning insights for possible use in COVID-19, Stem Cells Transl Med, 9 (2020) 1163–1173.

[9] B.P. Hurley, B.A. McCormick, Multiple roles of phospholipase A2 during lung infection and inflammation, Infect Immun, 76 (2008) 2259–2272.

[10] X. Yang, Y. Yu, J. Xu, H. Shu, J. Xia, H. Liu, Y. Wu, L. Zhang, Z. Yu, M. Fang, T. Yu, Y. Wang, S. Pan, X. Zou, S. Yuan, Y. Shang, Clinical course and outcomes of critically ill patients with SARS-CoV-2 pneumonia in Wuhan, China: a single-centered, retrospective, observational study, Lancet Respir Med, 8 (2020) 475–481.

[11] K.S.D. Kothapalli, H.G. Park, J.T. Brenna, Polyunsaturated fatty acid biosynthesis pathway and genetics. implications for interindividual variability in prothrombotic, inflammatory conditions such as COVID-19(,, bigstar, bigstar bigstar), Prostaglandins Leukot Essent Fatty Acids, 162 (2020) 102183.

[12] Worldometer., Countries where COVID-19 has spread, https://www.worldometers.info/coronavirus/countries-where-coronavirus-has-spread/, (2021).

[13] P.C. Calder, Omega-3 polyunsaturated fatty acids and inflammatory processes: nutrition or pharmacology?, Br J Clin Pharmacol, 75 (2013) 645–662.

[14] P.C. Norris, E.A. Dennis, Omega-3 fatty acids cause dramatic changes in TLR4 and purinergic eicosanoid signaling, Proc Natl Acad Sci U S A, 109 (2012) 8517–8522.

[15] J.K. Innes, P.C. Calder, Marine Omega-3 (N-3) Fatty Acids for Cardiovascular Health: An Update for 2020, Int J Mol Sci, 21 (2020).

[16] O.A. Byelashov, A.J. Sinclair, G. Kaur, Dietary sources, current intakes, and nutritional role of omega-3 docosapentaenoic acid, Lipid Technol, 27 (2015) 79–82.

[17] S. Ghasemi Fard, F. Wang, A.J. Sinclair, G. Elliott, G.M. Turchini, How does high DHA fish oil affect health? A systematic review of evidence, Crit Rev Food Sci Nutr, 59 (2019) 1684–1727.

[18] W. Wahli, L. Michalik, PPARs at the crossroads of lipid signaling and inflammation, Trends Endocrinol Metab, 23 (2012) 351–363.

[19] S.D. Clarke, The multi-dimensional regulation of gene expression by fatty acids: polyunsaturated fats as nutrient sensors, Curr Opin Lipidol, 15 (2004) 13–18.

[20] E.A. Dennis, P.C. Norris, Eicosanoid storm in infection and inflammation, Nat Rev Immunol, 15 (2015) 511–523.

[21] W.J. Park, K.S. Kothapalli, P. Lawrence, J.T. Brenna, FADS2 function loss at the cancer hotspot 11q13 locus diverts lipid signaling precursor synthesis to unusual eicosanoid fatty acids, PLoS One, 6 (2011) e28186.

[22] N. Morina, G. Bocari, A. Iljazi, K. Hyseini, G. Halac, Maximum Time of the Effect of Antileukotriene - Zileuton in Treatment of Patients with Bronchial Asthma, Acta Inform Med, 24 (2016) 16–19.

[23] J. P.J.H., A.A. Papamandjaris, Lipids: cellular metabolism., In: Erdman JW, Macdonald IA, Zeisel SH, eds. Present Knowledge in Nutrition. 10th ed. Washington, DC: Wiley-Blackwell., (2012) 132–148.

[24] P.J.H. Jones, T. Rideout, Lipids, sterols, and their metabolites., In: Ross AC, Caballero B, Cousins RJ, Tucker KL, Ziegler TR, eds. Modern Nutrition in Health and Disease. 11th ed. Baltimore, MD: Lippincott Williams & Wilkins., (2014).

[25] N. (ODS), Omega-3 Fatty Acids, https://ods.od.nih.gov/factsheets/Omega3FattyAcids-HealthProfessional/, (2021).

[26] H. Chappus-McCendie, L. Chevalier, C. Roberge, M. Plourde, Omega-3 PUFA metabolism and brain modifications during aging, Prog Neuropsychopharmacol Biol Psychiatry, 94 (2019) 109662.

[27] M. Ciriaco, P. Ventrice, G. Russo, M. Scicchitano, G. Mazzitello, F. Scicchitano, E. Russo, Corticosteroid-related central nervous system side effects, J Pharmacol Pharmacother, 4 (2013) S94–98.

[28] S.D. Glockler-Lauf, Y. Finkelstein, J. Zhu, L.Y. Feldman, T. To, Montelukast and Neuropsychiatric Events in Children with Asthma: A Nested Case-Control Study, J Pediatr, 209 (2019) 176–182 e174.

[29] R.H. Meyboom, N. de Graaf-Breederveld, Budesonide and psychic side effects, Ann Intern Med, 109 (1988) 683.

[30] L. Peyro-Saint-Paul, P. Besnier, L. Demessine, M. Biour, D. Hillaire-Buys, C. de Canecaude, S. Fedrizzi, J.J. Parienti, Cushing’s syndrome due to interaction between ritonavir or cobicistat and corticosteroids: a case-control study in the French Pharmacovigilance Database, J Antimicrob Chemother, 74 (2019) 3291–3294.

[31] C.M. Ajimura, N. Jagan, L.E. Morrow, M.A. Malesker, Drug Interactions With Oral Inhaled Medications, J Pharm Technol, 34 (2018) 273–280.

[32] F. Ferrau, F. Ceccato, S. Cannavo, C. Scaroni, What we have to know about corticosteroids use during Sars-Cov-2 infection, J Endocrinol Invest, 44 (2021) 693–701.

[33] A. Asher, N.L. Tintle, M. Myers, L. Lockshon, H. Bacareza, W.S. Harris, Blood omega-3 fatty acids and death from COVID-19: A pilot study, Prostaglandins Leukot Essent Fatty Acids, 166 (2021) 102250.

[34] S. Adams, A.L. Lopata, C.M. Smuts, R. Baatjies, M.F. Jeebhay, Relationship between Serum Omega-3 Fatty Acid and Asthma Endpoints, Int J Environ Res Public Health, 16 (2018).

[35] S.W. Njoroge, M. Laposata, W. Katrangi, A.C. Seegmiller, DHA and EPA reverse cystic fibrosis-related FA abnormalities by suppressing FA desaturase expression and activity, J Lipid Res, 53 (2012) 257–265.

[36] M. Parish, F. Valiyi, H. Hamishehkar, S. Sanaie, M. Asghari Jafarabadi, S.E. Golzari, A. Mahmoodpoor, The Effect of Omega-3 Fatty Acids on ARDS: A Randomized Double-Blind Study, Adv Pharm Bull, 4 (2014) 555–561.

[37] C. Lemoine, E. Brigham, H. Woo, A. Koch, C. Hanson, K. Romero, N. Putcha, M. McCormack, N. Hansel, Relationship between Omega-3 and Omega-6 Fatty Acid Intake and Chronic Obstructive Pulmonary Disease Morbidity, Ann Am Thorac Soc, 17 (2020) 378–381.

[38] B.S. van der Meij, J.A. Langius, E.F. Smit, M.D. Spreeuwenberg, B.M. von Blomberg, A.C. Heijboer, M.A. Paul, P.A. van Leeuwen, Oral nutritional supplements containing (n-3) polyunsaturated fatty acids affect the nutritional status of patients with stage III non-small cell lung cancer during multimodality treatment, J Nutr, 140 (2010) 1774–1780.

[39] H.A. Blair, S. Dhillon, Omega-3 carboxylic acids (Epanova): a review of its use in patients with severe hypertriglyceridemia, Am J Cardiovasc Drugs, 14 (2014) 393–400.

[40] L. Chevalier, M. Plourde, Comparison of pharmacokinetics of omega-3 fatty acid supplements in monoacylglycerol or ethyl ester in humans: a randomized controlled trial, Eur J Clin Nutr, 75 (2021) 680–688.

[41] L.A. Deinema, A.J. Vingrys, C.Y. Wong, D.C. Jackson, H.R. Chinnery, L.E. Downie, A Randomized, Double-Masked, Placebo-Controlled Clinical Trial of Two Forms of Omega-3 Supplements for Treating Dry Eye Disease, Ophthalmology, 124 (2017) 43–52.

[42] K.M. Doyle, D.A. Bird, S. al-Salihi, Y. Hallaq, J.E. Cluette-Brown, K.A. Goss, M. Laposata, Fatty acid ethyl esters are present in human serum after ethanol ingestion, J Lipid Res, 35 (1994) 428–437.

[43] P.C. Calder, Intravenous Lipid Emulsions to Deliver Bioactive Omega-3 Fatty Acids for Improved Patient Outcomes, Mar Drugs, 17 (2019).

[44] L. Anez-Bustillos, D.T. Dao, A.K. Potemkin, A.R. Perez-Atayde, B.P. Raphael, A.N. Carey, D.S. Kamin, J.R. Thiagarajah, M. Crowley, K.M. Gura, M. Puder, An Intravenous Fish Oil-Based Lipid Emulsion Successfully Treats Intractable Pruritus and Cholestasis in a Patient with Microvillous Inclusion Disease, Hepatology, 69 (2019) 1353–1356.

[45] D.S. Dzhalilova, A.M. Kosyreva, M.E. Diatroptov, E.A. Ponomarenko, I.S. Tsvetkov, N.A. Zolotova, V.A. Mkhitarov, D.N. Khochanskiy, O.V. Makarova, Dependence of the severity of the systemic inflammatory response on resistance to hypoxia in male Wistar rats, J Inflamm Res, 12 (2019) 73–86.

[46] S. Saadat, F. Beheshti, V.R. Askari, M. Hosseini, N. Mohamadian Roshan, M.H. Boskabady, Aminoguanidine affects systemic and lung inflammation induced by lipopolysaccharide in rats, Respir Res, 20 (2019) 96.

[47] A. Yokoyama, T. Hamazaki, A. Ohshita, N. Kohno, K. Sakai, G.D. Zhao, H. Katayama, K. Hiwada, Effect of aerosolized docosahexaenoic acid in a mouse model of atopic asthma, Int Arch Allergy Immunol, 123 (2000) 327–332.

[48] R. Renne, A. Brix, J. Harkema, R. Herbert, B. Kittel, D. Lewis, T. March, K. Nagano, M. Pino, S. Rittinghausen, M. Rosenbruch, P. Tellier, T. Wohrmann, Proliferative and nonproliferative lesions of the rat and mouse respiratory tract, Toxicol Pathol, 37 (2009) 5S–73S.

[49] L. Arbibe, K. Koumanov, D. Vial, C. Rougeot, G. Faure, N. Havet, S. Longacre, B.B. Vargaftig, G. Bereziat, D.R. Voelker, C. Wolf, L. Touqui, Generation of lyso-phospholipids from surfactant in acute lung injury is mediated by type-II phospholipase A2 and inhibited by a direct surfactant protein A-phospholipase A2 protein interaction, J Clin Invest, 102 (1998) 1152–1160.

[50] D. Vial, M. Senorale-Pose, N. Havet, L. Molio, B.B. Vargaftig, L. Touqui, Expression of the type-II phospholipase A2 in alveolar macrophages. Down-regulation by an inflammatory signal, J Biol Chem, 270 (1995) 17327–17332.

[51] M. Alaoui-El-Azher, Y. Wu, N. Havet, A. Israel, A. Lilienbaum, L. Touqui, Arachidonic acid differentially affects basal and lipopolysaccharide-induced sPLA(2)-IIA expression in alveolar macrophages through NF-kappaB and PPAR-gamma-dependent pathways, Mol Pharmacol, 61 (2002) 786–794.

[52] A.M. Lone, K. Tasken, Proinflammatory and immunoregulatory roles of eicosanoids in T cells, Front Immunol, 4 (2013) 130.

[53] H. Harizi, J.B. Corcuff, N. Gualde, Arachidonic-acid-derived eicosanoids: roles in biology and immunopathology, Trends Mol Med, 14 (2008) 461–469.

[54] C. Besenboeck, S. Cvitic, U. Lang, G. Desoye, C. Wadsack, Going into labor and beyond: phospholipase A2 in pregnancy, Reproduction, 151 (2016) R91–R102.

[55] M.F. McCarty, J.J. DiNicolantonio, Minimizing Membrane Arachidonic Acid Content as a Strategy for Controlling Cancer: A Review, Nutr Cancer, 70 (2018) 840–850.

[56] R. Wall, R.P. Ross, G.F. Fitzgerald, C. Stanton, Fatty acids from fish: the anti-inflammatory potential of long-chain omega-3 fatty acids, Nutr Rev, 68 (2010) 280–289.

[57] F. Norambuena, S. Morais, J.A. Emery, G.M. Turchini, Arachidonic Acid and Eicosapentaenoic Acid Metabolism in Juvenile Atlantic Salmon as Affected by Water Temperature, PLoS One, 10 (2015) e0143622.

[58] S.E. Wenzel, Arachidonic acid metabolites: mediators of inflammation in asthma, Pharmacotherapy, 17 (1997) 3S–12S.

[59] A.M. van der Does, M. Heijink, O.A. Mayboroda, L.J. Persson, M. Aanerud, P. Bakke, T.M. Eagan, P.S. Hiemstra, M. Giera, Dynamic differences in dietary polyunsaturated fatty acid metabolism in sputum of COPD patients and controls, Biochim Biophys Acta Mol Cell Biol Lipids, 1864 (2019) 224–233.

[60] J.S. Kim, B.T. Steffen, A.J. Podolanczuk, S.M. Kawut, I. Noth, G. Raghu, E.D. Michos, E.A. Hoffman, G.T. Axelsson, G. Gudmundsson, V. Gudnason, E.F. Gudmundsson, R.A. Murphy, J. Dupuis, H. Xu, R.S. Vasan, G.T. O’Connor, W.S. Harris, G.M. Hunninghake, R.G. Barr, M.Y. Tsai, D.J. Lederer, Associations of omega-3 Fatty Acids With Interstitial Lung Disease and Lung Imaging Abnormalities Among Adults, Am J Epidemiol, 190 (2021) 95–108.

[61] J. Allaire, P. Couture, M. Leclerc, A. Charest, J. Marin, M.C. Lepine, D. Talbot, A. Tchernof, B. Lamarche, A randomized, crossover, head-to-head comparison of eicosapentaenoic acid and docosahexaenoic acid supplementation to reduce inflammation markers in men and women: the Comparing EPA to DHA (ComparED) Study, Am J Clin Nutr, 104 (2016) 280–287.

[62] M.H. Liu, A.H. Lin, S.H. Lu, R.Y. Peng, T.S. Lee, Y.R. Kou, Eicosapentaenoic acid attenuates cigarette smoke-induced lung inflammation by inhibiting ROS-sensitive inflammatory signaling, Front Physiol, 5 (2014) 440.

[63] C. Morin, R. Hiram, E. Rousseau, P.U. Blier, S. Fortin, Docosapentaenoic acid monoacylglyceride reduces inflammation and vascular remodeling in experimental pulmonary hypertension, Am J Physiol Heart Circ Physiol, 307 (2014) H574–586.

[64] R. Adili, M. Hawley, M. Holinstat, Regulation of platelet function and thrombosis by omega-3 and omega-6 polyunsaturated fatty acids, Prostaglandins Other Lipid Mediat, 139 (2018) 10–18.

[65] J.M. Rudd, M. Tamil Selvan, S. Cowan, Y.F. Kao, C.C. Midkiff, S. Narayanan, A. Ramachandran, J.W. Ritchey, C.A. Miller, Clinical and Histopathologic Features of a Feline SARS-CoV-2 Infection Model Are Analogous to Acute COVID-19 in Humans, Viruses, 13 (2021).

[66] WHO, https://www.who.int/news-room/fact-sheets/detail/chronic-obstructive-pulmonary-disease-(copd), (2019).

[67] E. Bucchioni, S.A. Kharitonov, L. Allegra, P.J. Barnes, High levels of interleukin-6 in the exhaled breath condensate of patients with COPD, Respir Med, 97 (2003) 1299–1302.

[68] L.K. Lundblad, J. Thompson-Figueroa, T. Leclair, M.J. Sullivan, M.E. Poynter, C.G. Irvin, J.H. Bates, Tumor necrosis factor-alpha overexpression in lung disease: a single cause behind a complex phenotype, Am J Respir Crit Care Med, 171 (2005) 1363–1370.

[69] M. Konigshoff, N. Kneidinger, O. Eickelberg, TGF-beta signaling in COPD: deciphering genetic and cellular susceptibilities for future therapeutic regimen, Swiss Med Wkly, 139 (2009) 554–563.

[70] A.M. Gearhart, R. Cavallazzi, P. Peyrani, T.L. Wiemken, S. Furmanek, A. Reyes-Vega, U. Gauhar, H. Rivas-Perez, J. Roman, J.A. Ramirez, R. Fernandez-Botran, Lung cytokines and systemic inflammation in patients with COPD., Univ. Louisville J. Resp. Infect, 1 (2017) 13–18.

[71] Y. Zou, X. Chen, J. Liu, D.B. Zhou, X. Kuang, J. Xiao, Q. Yu, X. Lu, W. Li, B. Xie, Q. Chen, Serum IL-1beta and IL-17 levels in patients with COPD: associations with clinical parameters, Int J Chron Obstruct Pulmon Dis, 12 (2017) 1247–1254.

[72] J. Li, T. Lan, C. Zhang, C. Zeng, J. Hou, Z. Yang, M. Zhang, J. Liu, B. Liu, Reciprocal activation between IL-6/STAT3 and NOX4/Akt signalings promotes proliferation and survival of non-small cell lung cancer cells, Oncotarget, 6 (2015) 1031–1048.

[73] G.S. Shang, L. Liu, Y.W. Qin, IL-6 and TNF-alpha promote metastasis of lung cancer by inducing epithelial-mesenchymal transition, Oncol Lett, 13 (2017) 4657–4660.

[74] Y. Hasegawa, S. Takanashi, Y. Kanehira, T. Tsushima, T. Imai, K. Okumura, Transforming growth factor-beta1 level correlates with angiogenesis, tumor progression, and prognosis in patients with nonsmall cell lung carcinoma, Cancer, 91 (2001) 964–971.

[75] R. Wang, M. Lu, J. Zhang, S. Chen, X. Luo, Y. Qin, H. Chen, Increased IL-10 mRNA expression in tumor-associated macrophage correlated with late stage of lung cancer, J Exp Clin Cancer Res, 30 (2011) 62.

[76] E.B. Garon, J. Chih-Hsin Yang, S.M. Dubinett, The Role of Interleukin 1beta in the Pathogenesis of Lung Cancer, JTO Clin Res Rep, 1 (2020) 100001.

[77] S.K. Dhar, V. K, S. Damodar, S. Gujar, M. Das, IL-6 and IL-10 as predictors of disease severity in COVID-19 patients: results from meta-analysis and regression, Heliyon, 7 (2021) e06155.

[78] J.C. de Rivero Vaccari, W.D. Dietrich, R.W. Keane, J.P. de Rivero Vaccari, The Inflammasome in Times of COVID-19, Front Immunol, 11 (2020) 583373.

[79] L.C.K. Bell, C. Meydan, J. Kim, J. Foox, D. Butler, C.E. Mason, S.D. Shapira, M. Noursadeghi, G. Pollara, Transcriptional response modules characterize IL-1beta and IL-6 activity in COVID-19, iScience, 24 (2021) 101896.

[80] H. Han, Q. Ma, C. Li, R. Liu, L. Zhao, W. Wang, P. Zhang, X. Liu, G. Gao, F. Liu, Y. Jiang, X. Cheng, C. Zhu, Y. Xia, Profiling serum cytokines in COVID-19 patients reveals IL-6 and IL-10 are disease severity predictors, Emerg Microbes Infect, 9 (2020) 1123–1130.

[81] L. Lu, H. Zhang, D.J. Dauphars, Y.W. He, A Potential Role of Interleukin 10 in COVID-19 Pathogenesis, Trends Immunol, 42 (2021) 3–5.

[82] A. Copaescu, O. Smibert, A. Gibson, E.J. Phillips, J.A. Trubiano, The role of IL-6 and other mediators in the cytokine storm associated with SARS-CoV-2 infection, J Allergy Clin Immunol, 146 (2020) 518–534 e511.

[83] V. Azmy, K. Kaman, D. Tang, H. Zhao, C. Dela Cruz, J.E. Topal, M. Malinis, C.C. Price, Cytokine Profiles Before and After Immune Modulation in Hospitalized Patients with COVID-19, J Clin Immunol, 41 (2021) 738–747.

[84] B. Diao, C. Wang, Y. Tan, X. Chen, Y. Liu, L. Ning, L. Chen, M. Li, Y. Liu, G. Wang, Z. Yuan, Z. Feng, Y. Zhang, Y. Wu, Y. Chen, Reduction and Functional Exhaustion of T Cells in Patients With Coronavirus Disease 2019 (COVID-19), Front Immunol, 11 (2020) 827.

[85] D.M. Del Valle, S. Kim-Schulze, H.H. Huang, N.D. Beckmann, S. Nirenberg, B. Wang, Y. Lavin, T.H. Swartz, D. Madduri, A. Stock, T.U. Marron, H. Xie, M. Patel, K. Tuballes, O. Van Oekelen, A. Rahman, P. Kovatch, J.A. Aberg, E. Schadt, S. Jagannath, M. Mazumdar, A.W. Charney, A. Firpo-Betancourt, D.R. Mendu, J. Jhang, D. Reich, K. Sigel, C. Cordon-Cardo, M. Feldmann, S. Parekh, M. Merad, S. Gnjatic, An inflammatory cytokine signature predicts COVID-19 severity and survival, Nat Med, 26 (2020) 1636–1643.

[86] H.S. Vatansever, E. Becer, Relationship between IL-6 and COVID-19: to be considered during treatment, Future Virol, 15 (12) (2021) 817–822.

[87] M.C. Peters, K.W. McGrath, G.A. Hawkins, A.T. Hastie, B.D. Levy, E. Israel, B.R. Phillips, D.T. Mauger, S.A. Comhair, S.C. Erzurum, M.W. Johansson, N.N. Jarjour, A.M. Coverstone, M. Castro, F. Holguin, S.E. Wenzel, P.G. Woodruff, E.R. Bleecker, J.V. Fahy, L. National Heart, P. Blood Institute Severe Asthma Research, Plasma interleukin-6 concentrations, metabolic dysfunction, and asthma severity: a cross-sectional analysis of two cohorts, Lancet Respir Med, 4 (2016) 574–584.

[88] M. Rincon, C.G. Irvin, Role of IL-6 in asthma and other inflammatory pulmonary diseases, Int J Biol Sci, 8 (2012) 1281–1290.

[89] B.K. Itariu, M. Zeyda, E.E. Hochbrugger, A. Neuhofer, G. Prager, K. Schindler, A. Bohdjalian, D. Mascher, S. Vangala, M. Schranz, M. Krebs, M.G. Bischof, T.M. Stulnig, Long-chain n-3 PUFAs reduce adipose tissue and systemic inflammation in severely obese nondiabetic patients: a randomized controlled trial, Am J Clin Nutr, 96 (2012) 1137–1149.

[90] Y. Ma, C.E. Smith, C.Q. Lai, M.R. Irvin, L.D. Parnell, Y.C. Lee, L.D. Pham, S. Aslibekyan, S.A. Claas, M.Y. Tsai, I.B. Borecki, E.K. Kabagambe, J.M. Ordovas, D.M. Absher, D.K. Arnett, The effects of omega-3 polyunsaturated fatty acids and genetic variants on methylation levels of the interleukin-6 gene promoter, Mol Nutr Food Res, 60 (2016) 410–419.

[91] H. Zhao, Y. Chan-Li, S.L. Collins, Y. Zhang, R.W. Hallowell, W. Mitzner, M.R. Horton, Pulmonary delivery of docosahexaenoic acid mitigates bleomycin-induced pulmonary fibrosis, BMC Pulm Med, 14 (2014) 64.

[92] M.J. Page, J. Bester, E. Pretorius, The inflammatory effects of TNF-alpha and complement component 3 on coagulation, Sci Rep, 8 (2018) 1812.

[93] H. Akbari, R. Tabrizi, K.B. Lankarani, H. Aria, S. Vakili, F. Asadian, S. Noroozi, P. Keshavarz, S. Faramarz, The role of cytokine profile and lymphocyte subsets in the severity of coronavirus disease 2019 (COVID-19): A systematic review and meta-analysis, Life Sci, 258 (2020) 118167.

[94] C. Huang, Y. Wang, X. Li, L. Ren, J. Zhao, Y. Hu, L. Zhang, G. Fan, J. Xu, X. Gu, Z. Cheng, T. Yu, J. Xia, Y. Wei, W. Wu, X. Xie, W. Yin, H. Li, M. Liu, Y. Xiao, H. Gao, L. Guo, J. Xie, G. Wang, R. Jiang, Z. Gao, Q. Jin, J. Wang, B. Cao, Clinical features of patients infected with 2019 novel coronavirus in Wuhan, China, Lancet, 395 (2020) 497–506.

[95] S. Sanjabi, L.A. Zenewicz, M. Kamanaka, R.A. Flavell, Anti-inflammatory and pro-inflammatory roles of TGF-beta, IL-10, and IL-22 in immunity and autoimmunity, Curr Opin Pharmacol, 9 (2009) 447–453.

[96] T. Korn, E. Bettelli, M. Oukka, V.K. Kuchroo, IL-17 and Th17 Cells, Annu Rev Immunol, 27 (2009) 485–517.

[97] V. Dardalhon, A. Awasthi, H. Kwon, G. Galileos, W. Gao, R.A. Sobel, M. Mitsdoerffer, T.B. Strom, W. Elyaman, I.C. Ho, S. Khoury, M. Oukka, V.K. Kuchroo, IL-4 inhibits TGF-beta-induced Foxp3+ T cells and, together with TGF-beta, generates IL-9+ IL-10+ Foxp3(-) effector T cells, Nat Immunol, 9 (2008) 1347–1355.

[98] M. Veldhoen, C. Uyttenhove, J. van Snick, H. Helmby, A. Westendorf, J. Buer, B. Martin, C. Wilhelm, B. Stockinger, Transforming growth factor-beta ‘reprograms’ the differentiation of T helper 2 cells and promotes an interleukin 9-producing subset, Nat Immunol, 9 (2008) 1341–1346.

[99] K.M. P, K. Sivashanmugam, M. Kandasamy, R. Subbiah, V. Ravikumar, Repurposing of histone deacetylase inhibitors: A promising strategy to combat pulmonary fibrosis promoted by TGF-beta signalling in COVID-19 survivors, Life Sci, 266 (2021) 118883.

[100] Y. Xiong, Y. Liu, L. Cao, D. Wang, M. Guo, A. Jiang, D. Guo, W. Hu, J. Yang, Z. Tang, H. Wu, Y. Lin, M. Zhang, Q. Zhang, M. Shi, Y. Liu, Y. Zhou, K. Lan, Y. Chen, Transcriptomic characteristics of bronchoalveolar lavage fluid and peripheral blood mononuclear cells in COVID-19 patients, Emerg Microbes Infect, 9 (2020) 761–770.

[101] J.F. Pittet, M.J. Griffiths, T. Geiser, N. Kaminski, S.L. Dalton, X. Huang, L.A. Brown, P.J. Gotwals, V.E. Koteliansky, M.A. Matthay, D. Sheppard, TGF-beta is a critical mediator of acute lung injury, J Clin Invest, 107 (2001) 1537–1544.

[102] Y. Aschner, G.P. Downey, Transforming Growth Factor-beta: Master Regulator of the Respiratory System in Health and Disease, Am J Respir Cell Mol Biol, 54 (2016) 647–655.

[103] C.H. Maris, C.P. Chappell, J. Jacob, Interleukin-10 plays an early role in generating virus-specific T cell anergy, BMC Immunol, 8 (2007) 8.

[104] G. Lopez-Castejon, D. Brough, Understanding the mechanism of IL-1beta secretion, Cytokine Growth Factor Rev, 22 (2011) 189–195.

[105] A. Jain, R.A. Irizarry-Caro, M.M. McDaniel, A.S. Chawla, K.R. Carroll, G.R. Overcast, N.H. Philip, A. Oberst, A.V. Chervonsky, J.D. Katz, C. Pasare, T cells instruct myeloid cells to produce inflammasome-independent IL-1beta and cause autoimmunity, Nat Immunol, 21 (2020) 65–74.

[106] S. Baghaki, C.E. Yalcin, H.S. Baghaki, S.Y. Aydin, B. Daghan, E. Yavuz, COX2 inhibition in the treatment of COVID-19: Review of literature to propose repositioning of celecoxib for randomized controlled studies, Int J Infect Dis, 101 (2020) 29–32.

[107] L. Landi, C. Ravaglia, E. Russo, P. Cataleta, M. Fusari, A. Boschi, D. Giannarelli, F. Facondini, I. Valentini, I. Panzini, L. Lazzari-Agli, P. Bassi, E. Marchionni, R. Romagnoli, R. De Giovanni, M. Assirelli, F. Baldazzi, F. Pieraccini, G. Rametta, L. Rossi, L. Santini, I. Valenti, F. Cappuzzo, Blockage of interleukin-1beta with canakinumab in patients with Covid-19, Sci Rep, 10 (2020) 21775.

[108] A.R. Sousa, S.J. Lane, J.A. Nakhosteen, T.H. Lee, R.N. Poston, Expression of interleukin-1 beta (IL-1beta) and interleukin-1 receptor antagonist (IL-1ra) on asthmatic bronchial epithelium, Am J Respir Crit Care Med, 154 (1996) 1061–1066.

[109] A.I. Ritchie, A. Singanayagam, Immunosuppression for hyperinflammation in COVID-19: a double-edged sword?, Lancet, 395 (2020) 1111.

[110] A. Singanayagam, N. Glanville, J.L. Girkin, Y.M. Ching, A. Marcellini, J.D. Porter, M. Toussaint, R.P. Walton, L.J. Finney, J. Aniscenko, J. Zhu, M.B. Trujillo-Torralbo, M.A. Calderazzo, C. Grainge, S.L. Loo, P.C. Veerati, P.S. Pathinayake, K.S. Nichol, A.T. Reid, P.L. James, R. Solari, P.A.B. Wark, D.A. Knight, M.F. Moffatt, W.O. Cookson, M.R. Edwards, P. Mallia, N.W. Bartlett, S.L. Johnston, Corticosteroid suppression of antiviral immunity increases bacterial loads and mucus production in COPD exacerbations, Nat Commun, 9 (2018) 2229.

[111] B.J. Thomas, R.A. Porritt, P.J. Hertzog, P.G. Bardin, M.D. Tate, Glucocorticosteroids enhance replication of respiratory viruses: effect of adjuvant interferon, Sci Rep, 4 (2014) 7176.

[112] C.D. Russell, J.E. Millar, J.K. Baillie, Clinical evidence does not support corticosteroid treatment for 2019-nCoV lung injury, Lancet, 395 (2020) 473–475.

[113] P. Mehta, D.F. McAuley, M. Brown, E. Sanchez, R.S. Tattersall, J.J. Manson, U.K. Hlh Across Speciality Collaboration, COVID-19: consider cytokine storm syndromes and immunosuppression, Lancet, 395 (2020) 1033–1034.

[114] B.E. Kula, J. Cornelius, M. Clancy, H. Nguyen, I.S. Schwartz, Invasive Mould Disease in Fatal COVID-19: A Systematic Review of Autopsies medRxiv, (2021).

[115] A. Moorthy, R. Gaikwad, S. Krishna, R. Hegde, K.K. Tripathi, P.G. Kale, P.S. Rao, D. Haldipur, K. Bonanthaya, SARS-CoV-2, Uncontrolled Diabetes and Corticosteroids-An Unholy Trinity in Invasive Fungal Infections of the Maxillofacial Region? A Retrospective, Multi-centric Analysis, J Maxillofac Oral Surg, (2021) 1–8.

[116] G.R. Thompson Iii, O.A. Cornely, P.G. Pappas, T.F. Patterson, M. Hoenigl, J.D. Jenks, C.J. Clancy, M.H. Nguyen, Invasive Aspergillosis as an Under-recognized Superinfection in COVID-19, Open Forum Infect Dis, 7 (2020) ofaa242.

[117] D. Armstrong-James, J. Youngs, T. Bicanic, A. Abdolrasouli, D.W. Denning, E. Johnson, V. Mehra, T. Pagliuca, B. Patel, J. Rhodes, S. Schelenz, A. Shah, F.L. van de Veerdonk, P.E. Verweij, P.L. White, M.C. Fisher, Confronting and mitigating the risk of COVID-19 associated pulmonary aspergillosis, Eur Respir J, 56 (2020).

[118] M. Sen, S. Lahane, T.P. Lahane, R. Parekh, S.G. Honavar, Mucor in a Viral Land: A Tale of Two Pathogens, Indian J Ophthalmol, 69 (2021) 244–252.

[119] V.E. de Meijer, K.M. Gura, H.D. Le, J.A. Meisel, M. Puder, Fish oil-based lipid emulsions prevent and reverse parenteral nutrition-associated liver disease: the Boston experience, JPEN J Parenter Enteral Nutr, 33 (2009) 541–547.

[120] K.L. Calkins, M. Puder, K. Gura, The evolving use of intravenous lipid emulsions in the neonatal intensive care unit, Semin Perinatol, 43 (2019) 151155.

[121] K.M. Gura, M.H. Premkumar, K.L. Calkins, M. Puder, Fish Oil Emulsion Reduces Liver Injury and Liver Transplantation in Children with Intestinal Failure-Associated Liver Disease: A Multicenter Integrated Study, J Pediatr, 230 (2021) 46–54 e42.

[122] A.S. Abdelhamid, T.J. Brown, J.S. Brainard, P. Biswas, G.C. Thorpe, H.J. Moore, K.H. Deane, C.D. Summerbell, H.V. Worthington, F. Song, L. Hooper, Omega-3 fatty acids for the primary and secondary prevention of cardiovascular disease, Cochrane Database Syst Rev, 3 (2020) CD003177.

[123] C.J. Glueck, N. Khan, M. Riaz, J. Padda, Z. Khan, P. Wang, Titrating lovaza from 4 to 8 to 12 grams/day in patients with primary hypertriglyceridemia who had triglyceride levels >500 mg/dl despite conventional triglyceride lowering therapy, Lipids Health Dis, 11 (2012) 143.

[124] S. Doaei, S. Gholami, S. Rastgoo, M. Gholamalizadeh, F. Bourbour, S.E. Bagheri, F. Samipoor, M.E. Akbari, M. Shadnoush, F. Ghorat, S.A. Mosavi Jarrahi, N. Ashouri Mirsadeghi, A. Hajipour, P. Joola, A. Moslem, M.O. Goodarzi, The effect of omega-3 fatty acid supplementation on clinical and biochemical parameters of critically ill patients with COVID-19: a randomized clinical trial, J Transl Med, 19 (2021) 128.

[125] A.B. Mohabbat, N.M.L. Mohabbat, E.C. Wight, Fibromyalgia and Chronic Fatigue Syndrome in the Age of COVID-19, Mayo Clin Proc Innov Qual Outcomes, 4 (2020) 764–766.

[126] A.L. Komaroff, The tragedy of long COVID, https://www.health.harvard.edu/blog/the-tragedy-of-the-post-covid-long-haulers-202010152479, (2021).

[127] F. Salaffi, V. Giorgi, S. Sirotti, S. Bongiovanni, S. Farah, L. Bazzichi, D. Marotto, F. Atzeni, M. Rizzi, A. Batticciotto, G. Lombardi, M. Galli, P. Sarzi-Puttini, The effect of novel coronavirus disease-2019 (COVID-19) on fibromyalgia syndrome, Clin Exp Rheumatol, 39 Suppl 130 (2021) 72–77.

[128] S. Ozgocmen, S.A. Catal, O. Ardicoglu, A. Kamanli, Effect of omega-3 fatty acids in the management of fibromyalgia syndrome, Int J Clin Pharmacol Ther, 38 (2000) 362–363.

[129] G.D. Ko, N.B. Nowacki, L. Arseneau, M. Eitel, A. Hum, Omega-3 fatty acids for neuropathic pain: case series, Clin J Pain, 26 (2010) 168–172.

[130] I. Kostoglou-Athanassiou, L. Athanassiou, P. Athanassiou, The Effect of Omega-3 Fatty Acids on Rheumatoid Arthritis, Mediterr J Rheumatol, 31 (2020) 190–194.

[131] V. Wiwanitkit, Delirium, sleep, COVID-19 and melatonin, Sleep Med, 75 (2020) 542.

[132] E. Zambrelli, M. Canevini, O. Gambini, A. D’Agostino, Delirium and sleep disturbances in COVID-19: a possible role for melatonin in hospitalized patients?, Sleep Med, 70 (2020) 111.

[133] R. Zhang, X. Wang, L. Ni, X. Di, B. Ma, S. Niu, C. Liu, R.J. Reiter, COVID-19: Melatonin as a potential adjuvant treatment, Life Sci, 250 (2020) 117583.

[134] A. Shneider, A. Kudriavtsev, A. Vakhrusheva, Can melatonin reduce the severity of COVID-19 pandemic?, Int Rev Immunol, 39 (2020) 153–162.

[135] Y. Zhou, Y. Hou, J. Shen, R. Mehra, A. Kallianpur, D.A. Culver, M.U. Gack, S. Farha, J. Zein, S. Comhair, C. Fiocchi, T. Stappenbeck, T. Chan, C. Eng, J.U. Jung, L. Jehi, S. Erzurum, F. Cheng, A network medicine approach to investigation and population-based validation of disease manifestations and drug repurposing for COVID-19, PLoS Biol, 18 (2020) e3000970.

[136] D.R. McDougle, J.E. Watson, A.A. Abdeen, R. Adili, M.P. Caputo, J.E. Krapf, R.W. Johnson, K.A. Kilian, M. Holinstat, A. Das, Anti-inflammatory omega-3 endocannabinoid epoxides, Proc Natl Acad Sci U S A, 114 (2017) E6034–E6043.

[137] J.E. Watson, J.S. Kim, A. Das, Emerging class of omega-3 fatty acid endocannabinoids & their derivatives, Prostaglandins Other Lipid Mediat, 143 (2019) 106337.

[138] G. Esposito, M. Pesce, L. Seguella, W. Sanseverino, J. Lu, C. Corpetti, G. Sarnelli, The potential of cannabidiol in the COVID-19 pandemic, Br J Pharmacol, 177 (2020) 4967–4970.

[139] L.C. Nguyen, D. Yang, V. Nicolaescu, T.J. Best, T. Ohtsuki, S.N. Chen, J.B. Friesen, N. Drayman, A. Mohamed, C. Dann, D. Silva, H. Gula, K.A. Jones, J.M. Millis, B.C. Dickinson, S. Tay, S.A. Oakes, G.F. Pauli, D.O. Meltzer, G. Randall, M.R. Rosner, Cannabidiol Inhibits SARS-CoV-2 Replication and Promotes the Host Innate Immune Response, bioRxiv, (2021).

